# *Plasmodium vivax* relapse reservoirs in Ethiopia: microhaplotype sequencing informs recurrent infection origins

**DOI:** 10.64898/2026.11.12.765293

**Authors:** Ashley A. Osborne, Shazia Ruybal-Pesantez, Anjana Rai, Hidayat Trimarsanto, Kian Soon Hoon, Kim Piera, Mariana Kleinecke, Angela Rumaseb, Edwin Sutanto, Kamala Thriemer, Tamiru S. Degaga, Jimee Hwang, Ashenafi Assefa, Adugna Woyessa, Asrat Hailu, Aimee Taylor, Nicholas P. Day, Ric N. Price, Sarah Auburn

## Abstract

Determining the origin of recurrent *Plasmodium vivax* infections is critical for quantifying the public health burden of the hypnozoite reservoir, understanding transmission potential, and evaluating anti-relapse antimalarial efficacy. In Ethiopia, *P. vivax* is a major contributor to malaria burden, but the relative contribution of relapse to recurrent infections remains poorly quantified. We performed microhaplotype genotyping using a panel of 93 genome-wide markers and applied a probabilistic recurrence classification framework to recurrent *P. vivax* samples collected in two Ethiopian radical cure trials (IMPROV and OPRA). Genotyping was conducted on parasite isolates collected at enrolment and subsequent recurrences, with participants monitored over a 12-month follow-up period (IMPROV: 604 samples from 444 participants; OPRA: 621 samples from 342 participants). Recurrent infections were classified using a Bayesian hierarchical model implemented in *Pv3Rs*, which estimated marginal posterior probabilities of relapse, reinfection, or recrudescence based on genotyping data and background population dynamics. Priors were structured across biologically motivated time windows reflecting the natural history of *P. vivax* infection in Ethiopia and anticipated drug efficacy. Recurrences with a probability of relapse (p(L)) ≥ 0.75 were classified as relapses.

In the absence of primaquine, probable relapse was the dominant classification for primary recurrent infections in both trials (75.5%; 102/135) and for consecutive recurrent infections within individuals (75.2%; 204/271). Primaquine substantially reduced the proportion of participants experiencing recurrent infections attributable to probable relapse, from 57.3% to 1.7% in IMPROV and from 60.2% to 9.5% in OPRA. Polyclonal infection at enrolment was independently associated with increased odds of recurrence and a higher rate of probable relapse in OPRA, with a consistent but non-significant trend observed in IMPROV. These findings provide direct molecular evidence that the hypnozoite reservoir is the dominant driver of recurrent *P. vivax* infections in Ethiopia in the absence of radical cure. The marked reduction in probable relapse following primaquine treatment highlights the critical importance of expanding access to safe and effective radical cure strategies in Ethiopia and other comparable high-transmission settings.

## INTRODUCTION

*Plasmodium vivax* is an incredibly difficult human malaria species to eliminate due to its ability to form dormant liver-stage hypnozoites following an initial infection [1]. These hypnozoites can reactivate weeks to months after an initial infection, resulting in recurrent episodes of malaria (relapses), that are capable of sustaining recurrent illness episodes and ongoing transmission of the parasite [2]. Recurrent infections caused by relapses are clinically indistinguishable from reinfections caused by new mosquito inoculations, or recrudescence, resulting from a blood-stage treatment failure [3,4]. In most endemic settings, relapses are estimated to substantially outnumber reinfections, with some suggesting that relapses account for up to 80% of recurrent infections in individuals who are not treated with radical cure [5].

Determining the origin of recurrent *P. vivax* infections is critical for quantifying the true public health burden of the hypnozoite reservoir, including estimates surrounding transmission potential, and the evaluation of anti-relapse antimalarial efficacy regimens [6,7]. Accurately distinguishing relapses from reinfections and recrudescence, however, is methodologically challenging [6]. Recent advances in sequencing and analytical methodologies have begun to unravel the complex infection dynamics in high transmission settings [7]. Comparative genetics of paired parasite isolates collected at the time a patient is initially infected and enrolled into a clinical trial, and again at the time of subsequent parasite recurrence, can generate important information to inform whether recurrences are identical or highly related (e.g. siblings or half-siblings), or if they originate from unrelated reinfections [8]. Parasite relatedness can be quantified by estimating identity-by-descent (IBD), with recently developed software enabling characterisations from polyclonal infections [9]. Estimating the proportion of recurrences that are attributable to relapses cannot, however, be defined solely as pairs that exhibit high pairwise relatedness as IBD estimates are confounded by the complex genetics of sibling parasites and liver reservoirs. To address these limitations, probabilistic analytical frameworks have been developed that incorporate estimate of parasite relatedness and within-host multiplicity of infection within a Bayesian hierarchical structure, from which probabilities of whether a recurrent infection is due to relapse, reinfection, and recrudescence can be derived [10,11].

Genome-wide data from whole genome sequencing (WGS), is ideal for quantifying IBD, but is not feasible in many malaria-endemic settings. Initial studies attempted to address the relapse dilemma using genotyping data derived from 8-11 multiallelic markers, but statistical evaluations suggest that this approach was insufficient to achieve low errors in IBD [10,12]. Precision of IBD estimates can be improved substantially by using a larger panel of multiallelic markers distributed across the *P. vivax* genome. Amplicon-based sequencing of 93 microhaplotype markers, using highly parallel sequencing platforms, offers a cost effective and logistically efficient means to capture IBD accurately [13–16]. Microhaplotypes offer substantially greater discriminatory power to estimate IBD than single nucleotide polymorphisms (SNPs) on their own, while remaining more manageable at scale than WGS. This has led to them being increasingly adopted by research teams and national malaria control programmes (NMCPs) [13,14,17]. These advances in sequencing and analytical methodologies are instrumental in determining the contribution of relapse to the burden of *P. vivax* in high transmission regions. This will ultimately be critical to malaria elimination efforts and shrinking the geographic coverage of malaria worldwide [7].

In the Horn of Africa, *P. vivax* is a major cause of malaria. In 2024, Ethiopia accounted for approximately 97% of the *P. vivax* burden in Africa, with *P. vivax* infections becoming more prevalent in areas of the country that were previously dominated by *P. falciparum* [1,18]. In most endemic countries the recommended treatment for *P. vivax* malaria is a 3 day regimen of chloroquine, or an artemisinin-based combination therapy (ACT), combined with primaquine, usually administered at a total dose of 3.5mg/kg over 14 days [1,7]. The combination of blood-stage antimalarial treatment drugs with schizonticidal and hypnozoitocidal activity is known as radical cure. Blood-stage treatment of *P. vivax* with chloroquine appears to be highly effective in Ethiopia. There has been limited evidence of recrudescence reported, with treatment failures largely believed to be attributable to drug mal-absorption, poor drug quality, or re-infection [19,20]. Hesitancy when prescribing primaquine persists in Ethiopia. This is due to concerns about the risk of severe haemolysis in individuals with glucose-6-phosphate dehydrogenase (G6PD) deficiency and the limited availability of point-of-care G6PD testing [18–21].

Two large clinical trials have been conducted to determine the efficacy of radical cure regimens in Ethiopia and provide a unique opportunity to quantify the relative burden of relapse versus reinfection in high transmission regions of Ethiopia. The “Improving the radical cure of vivax malaria” (IMPROV) and “OPtimising the RAdical cure of vivax malaria” (OPRA) clinical trials were randomised controlled trials designed to evaluate the efficacy of different primaquine-based radical cure regimens against recurrent *P. vivax* relative to treatment of the blood-stage infection alone [19,21,22]. The primaquine-free arms provide a rare opportunity to observe natural recurrence dynamics directly. The trials were conducted in geographically and temporally distinct populations within Ethiopia, providing additional baseline population genetic data from historically relevant transmission periods on Ethiopia’s road to malaria elimination [19,21,23].

We applied microhaplotype genotyping and probabilistic recurrence classification methodologies to *P. vivax* parasite isolates collected from trial participants with uncomplicated *P. vivax* malaria, from the time of enrolment and for each subsequent recurrence over the course of a 12-month follow-up period, to quantify relapse and reinfection dynamics in this setting. We characterised baseline parasite population diversity and structure at each trial site prior to integrating within-host relatedness metrics and probabilistic classification methods to quantify the relative contribution of relapse to recurrent infections. Using these methods, we were able to assess whether infection complexity at enrolment was associated with subsequent risk of relapse, as well as the timing and frequency of relapse burden experienced over extended follow-up. Our analyses provide a molecular basis for reassessing the contribution of the hypnozoite reservoir to recurrent *P. vivax* in Ethiopia, with implications for radical cure policy in this and comparable endemic settings.

## METHODS

### Study Populations

The OPRA clinical trial (ClinicalTrials.gov NCT01680406) was conducted between 2012 and 2014 at two sites, Batu and Bishoftu, in central Ethiopia (**Supplementary Figure S1**) [19]. The four treatment arms, including artemether-lumefantrine (AL), artemether-lumefantrine plus primaquine (ALPQ), chloroquine (CQ), and chloroquine plus primaquine (CQPQ). Individuals in the two primaquine (PQ) arms received a standard low-dose course of PQ (total dose of 3.5 mg/kg administered over 14 days). PQ administration was semi-supervised during the initial episode, with directly observed treatment on days 2, 3, 7, 10, and 14; doses in between were self-administered. For all subsequent episodes of malaria, schizonticidal, as well as PQ treatment, was unsupervised. Individuals were followed for 12 months, and all recurrent episodes of malaria were treated during this period. The rate ratio of recurrent *P. vivax* parasitaemia was 5.1 to 6.4-fold lower in patients who were treated with PQ.

The IMPROV trial was a multicentre, randomised, double-blinded, placebo-controlled trial (ClinicalTrials.gov NCT01814683) conducted between 2017 and 2018 in Arba Minch and Metehara (**Supplementary Figure S1**) [21,23]. Individuals with microscopy-confirmed *P. vivax* malaria and normal G6PD activity were randomised to one of three treatment arms: CQ plus placebo (control), CQ plus a high-dose PQ (total dose 7mg/kg) administered either over 7 or 14 days. Every dose of PQ treatment was supervised. Participants were followed daily during treatment, then weekly for 8 weeks, and thereafter monthly for a total of 12 months. In this trial, the rate ratio of recurrent *P. vivax* infection was 5.3-fold lower in patients who were treated with high-dose PQ.

The risk of recurrence in the two primaquine arms in OPRA and the two primaquine arms in IMPROV were similar [19,21]. Hence for downstream analyses, the two primaquine arms in OPRA (ALPQ and CQPQ) were pooled into a single “PQ” group and compared to either the AL or CQ control arms. For IMPROV, the 7-day and 14-day PQ arms were pooled and compared to the CQ control arm [21]. These groupings were only applied to reporting and summary statistics. Genetic classification, including allele frequency estimation, background relatedness, and Pv3Rs posterior computations, were performed independently per trial, to accurately account for trial-specific temporal and geographic differences, and was unaffected by this reported grouping.

### Sample preparation and sequencing

A total of 363 individuals were genotyped from the OPRA trial (AL, N = 90; ALPQ, N = 79; CQ, N = 98; CQPQ, N = 96), and 453 from the IMPROV trial (Control, N = 94; 7-day PQ, N = 182; 14-day PQ, N = 177). Genotyping was performed using the vivaxGEN *P. vivax* microhaplotype panel, comprising 93 amplicons distributed across the *P. vivax* genome (PvP01v2 reference) [13,24]. Library preparation of the 93-plex marker panel was conducted using a single multiplexed primer amplification with pre-balanced, customised *rhAmpSeq* primers (*Integrated DNA Technologies*), followed by a second PCR to incorporate sample indexes and *Illumina* sequencing adaptors. Indexed products were pooled, with up to 384 samples per pool, bead-purified, prior to being quality-assessed for fragment size and concentration ahead of sequencing. Sequencing was performed with 150 bp paired-end reads on an *Illumina MiSeq* platform. Full assay design and validation are described in Kleinecke *et al.* 2025, with protocol details also available on protocols.io [13,24].

### Bioinformatic processing

The vivaxGEN *P. vivax* microhaplotype pipeline (github.com/vivaxgen/MicroHaps) was used to process genotyping data. Read quality trimming and adapter removal were performed using *fastp*, while amplicon sequence variants (ASVs) were called using the denoising *DADA2* software and custom postprocessing scripts [25,26]. For downstream analyses, indel variants were not included and loci with less than 25 read pairs per sample, were removed. To minimise the influence of genotyping error on allele frequency estimates, alleles present at less than 5% proportion relative to the major allele at a given locus, within a sample, were removed. Alleles required a minimum of 5 read pairs within a sample, and alleles detected in fewer than 5 samples overall were excluded. Samples were retained if at least 75 of the original 93 loci (∼80%) remained following these filters.

### Genetic measures of within-host diversity and population-level dynamics

The multiplicity of infection (MOI) for each isolate was estimated using the R-package *MOIRE,* a Bayesian method that estimates the effective MOI (eMOI) as the posterior mean number of co-infecting clones [27]. *MOIRE* was run using 200 burn-in iterations, 10,000 samples per chain, and 40 parallel tempering chains. To ensure MOIRE eMOI estimates were representative of anticipated within-population levels of diversity, we compared eMOI estimates to the mean number of alleles per loci in a given sample, as well as the naive complexity of infection (COI), or the max number of alleles at any given filtered loci. The mean eMOI was used as the primary measure of COI in downstream analyses to classify samples as monoclonal (eMOI ≤ 1.1) or polyclonal (eMOI > 1.1). An eMOI equal to 1 was not used to define “monoclonal” as posterior uncertainty produces mean values slightly above 1 that remain consistent with single-clone infections.

Pairwise identity-by-descent (IBD) between Day 0 samples, and between within-patient infection pairs, was estimated using the R package *Dcifer* [9]. The *Dcifer* package calculates genetic distance between both monoclonal and polyclonal infections by estimating relatedness from multiallelic genotyping data.

Baseline relatedness within each trial was estimated using Day 0 samples, to characterise background population connectivity prior to within-host analysis and to generate population-level allele frequencies (**Supplementary Results S1**). Enrolment was rolling in both trials, but a Mantel test confirmed that pairwise genetic distance was not confounded by enrolment timing when using months as the time interval [28]. Enrolment timing explained less than 0.4% and 0.11% of variance in pairwise relatedness in IMPROV (N = 433; r = 0.067; r² = 0.004; p = 0.0001) and OPRA (N = 328; r = 0.033; r² = 0.0011; p = 0.0001), respectively.

Highly related networks (IBD ≥ 25%) among Day 0 samples were driven predominantly by polyclonal infections and substantially reduced when restricted to monoclonal samples, confirming that polyclonality inflates background relatedness (**Supplementary Results S1**). This 25% threshold was further supported by the empirical distribution of relatedness among independent, cross-individual pairs of isolates collected at enrolment (Day 0). Overall, 98.6% of same-site pairs exhibited an IBD < 25% (N = 100,197 pairings), rising to 99.4% restricted to monoclonal pairs (N = 35,779). Differences in pairwise IBD across treatment arms were assessed using the Kruskal-Wallis test and pairwise Mann-Whitney U tests. Distributions are reported as mean and interquartile range (IQR = 25^th^ to 75th percentile) throughout.

### Recurrence classification framework

Probabilistic classification of recurrent infections was performed using the R-package *Pv3Rs*, which implements a Bayesian hierarchical model returning marginal posterior probabilities of relapse (L), reinfection (I), and recrudescence (C) across all within-individual infection episodes [10,11]. *Pv3Rs* explicitly models within-host multiplicity of infection which accounts for the probability that allele sharing between polyclonal infections arises by chance rather than by direct genetic descent. This is essential in high-transmission settings where polyclonal infections are common.

Priors were structured across three biologically motivated time windows rather than applied uniformly across the enrolment period, reflecting the anticipated natural history of *P. vivax* infection in Ethiopia and known drug efficacy (**Supplementary Table S1**).

1. The reinfection prior (pI) was derived as ***pI = 1 – exp(-λt)***; with ***λ = 0.002/day***, representing a continuous daily probability of acquiring a new infection that accumulates over time. This λ corresponds to a somewhat higher annual parasite incidence (API) than was recorded at the trial sites during the study period which should produce conservative lower-bound estimates of relapse, rather than inflating estimates, and is equivalent to a cumulative probability of acquiring *P. vivax* by day 90 at 17%, approximately [29].
2. The relapse prior (pL) was discounted in windows 2 and 3 by a factor of 0.19 for any of the primaquine treatment arms. This was derived from the 5.3-fold reduction in recurrence observed in the IMPROV clinical trial primaquine arms. This was consistent within the OPRA trial as well, which observed a 5.1-fold to 6.4-fold reduction in recurrence [19,21].
3. The recrudescence prior (pC) for window 1 was informed by the reported 8% late treatment failure in OPRA and the absence of documentation suggesting any *P. vivax* resistance to AL or CQ in Ethiopia [19,20]. AL clears substantially faster than CQ as lumefantrine has an average half-life of 3 to 6 days, upwards of 10 days in some patients. Window boundaries were set independently for AL arms (window 1 = ≤14 days; window 2 = 15 to 60 days) and CQ arms (window 1 = ≤35 days; window 2 = 36 to 120 days). A corresponding pharmacologically plausible threshold for post-hoc reassignment of raw recrudescence calls was set at 14 days for AL arms and 45 days for CQ arms.

**Supplementary Table 1:**
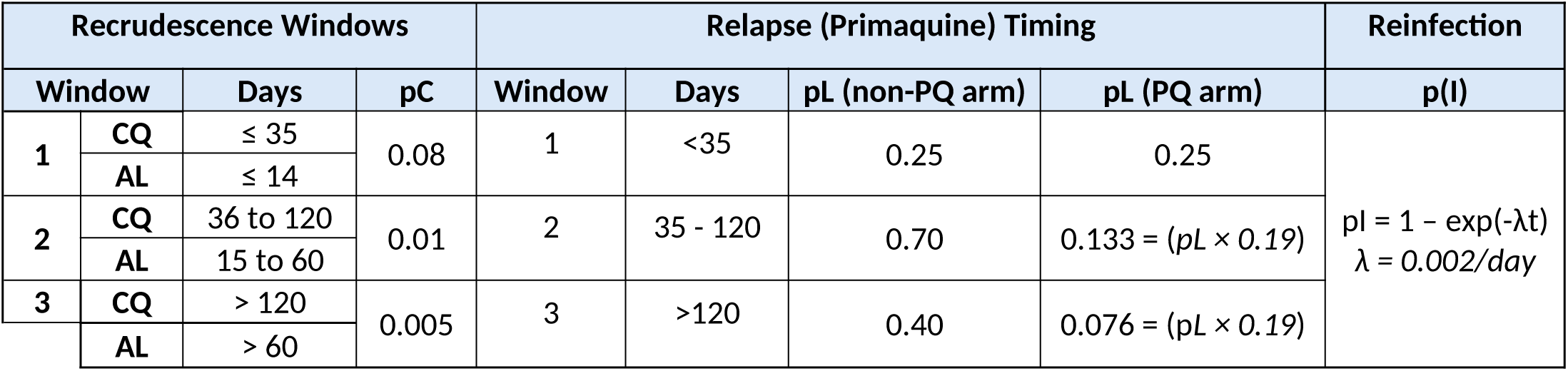
*Pv3rs* priors given Ethiopia’s reported *P. vivax* transmission intensity and dynamics of anti-malarial and anti-relapse drug pharmacologies in IMPROV and OPRA.

The interval used to assess pharmacological plausibility of a recrudescence was defined as the number of days between each pair’s sampled time points (i.e., time since the immediately preceding infection episode for that patient). This reflects that patients were treated at every observed recurrence in both trials, resetting the pharmacological clock at each treatment event. Recrudescence was rarely the most probable *Pv3Rs* classification for recurrence pairs in either trial. Given the residual ambiguity in these estimates and the absence of a tailored time-to-event model, recrudescence was not retained as a primary classification category in reported outputs but is reported in **Supplementary Table S2**.

Infection pairs were assessed under two analytical frameworks. First, only the sample at time of enrolment (D0) and the first recurrent infection (DR1) per individual were analysed, providing one independent observation per participant in which subsequent recurrences cannot influence *Pv3Rs* outputs. Second, all consecutive within-patient pairs were analysed jointly, comparing each recurrent infection to the immediately preceding infection (D0→DR1, DR1→DR2, DR2→DR3, etc.). An individual’s samples were included in pairwise analysis only up to, and including, the longest unbroken sequence of recurrences. This required a successfully genotyped sample on Day 0. Patients without a genotyped Day 0 sample were excluded entirely from all pairwise analyses. Where a patient’s recurrence sequence of events contained a gap (e.g. a recurrence event with no corresponding genotyped sample), only samples preceding that gap were retained for pairwise classification. Any samples following the gap were excluded as a pairwise comparison would then combine two, non-adjacent episodes, as though they were consecutive and not affected by the intermittent treatment administered to the patient. A total of 32 individuals were excluded from pairwise analysis under this rule, including 8 individuals from IMPROV who did not have a genotyped Day 0 sample. A further 23 individuals from IMPROV and 1 individual from OPRA were missing genotyping data for crucial recurrences. These individuals were included in the total genotyped-recurrence counts reported in the results, but are absent from all consecutive-pair, classification, and arm-level analyses thereafter.

*Pv3Rs* imposes a computational limit of cumulative MOI, with a maximum of 8 distinct clones across all modelled within-host episodes [11]. When this limit was exceeded, a stepwise fallback analysed each consecutive pair independently within the same time-window prior structure. Single pairs where MOI exceeded 8 could not be analysed. To assess robustness against within-host complexity, monoclonal-only pairs (both samples eMOI ≤ 1.1) were compared directly against polyclonal-involving pairs (at least one sample eMOI > 1.1) as two independent groups.

*Pv3Rs* assumes that all parasites within an infection episode draw from exactly two parental gametes. When this assumption is violated, including instances where half-sibling parasites draw from three or more parental alleles, the model can erroneously produce elevated posterior probabilities of reinfection p(I). These instances appear as outliers with high pairwise IBD, but unexpectedly low p(L). The number of affected pairs is anticipated to be small and study-level classification proportions should be robust to such individual-level discrepancies [30]. To mitigate erroneous and uncertain posterior probabilities, recurrence episodes were only classified as relapses if the probability of relapse p(L) was ≥ 0.75 and were classified as a reinfection if the p(L) was ≤ 0.25. Pairs with values between 0.25 and 0.75 were classified as indeterminate, to prevent classifying pairs with uncertain probabilities, or pairs that closely resemble multiple states of recurrence, in the final analysis.

The directionality of polyclonality was recorded for each classification pair. This could be: (1) A polyclonal Day 0 infection with a monoclonal recurrent infection pair (D0 to DR1), consistent with a single hypnozoite clone seeding a relapse infection; (2) A monoclonal Day 0 and a polyclonal recurrence, suggesting either a new polyclonal infection at the time of recurrence or a new infection alongside a relapse; (3) both polyclonal; or (4) both monoclonal [3,31]. This context was retained as a covariate for interpreting discordant classifications between software and relatedness estimates.

Batu and Bishoftu (OPRA) were analysed jointly using combined Day 0 allele frequencies, supported by the absence of a practically meaningful difference in background relatedness between the sites. From IMPROV, only Arba Minch was carried forward for classification analyses, since there was significantly higher polyclonality at Arba Minch relative to Metehara and an insufficient number of Metehara samples (N = 6 recurrence pairs) for estimating allele frequency reliably.

### Association between Day 0 clonality and the risk of recurrence or relapse

To assess whether Day 0 infection complexity was associated with an increased risk of subsequent recurrence or relapse, analyses were restricted to non-primaquine control arms (i.e. CQ alone in IMPROV; the AL and CQ arms in OPRA), where the hypnozoite reservoir was not pharmacologically cleared, and fitted separately per trial. This approach was designed to account for the different baseline polyclonality between IMPROV and OPRA. To estimate the association between Day 0 clonality and each parasitological outcome, after adjusting for treatment level differences in baseline recurrence rates, a logistic regression model was applied to all non-PQ individuals from both trials. This included an individual’s recurrence status (i.e. did or did not recur), Day 0 clonality, treatment arm, and probable relapses (p[L] ≥ 0.75). Recurrence was defined as any recorded recurrent sample, regardless of whether that sample was itself successfully genotyped. For the relapse outcome, only individuals whose recurrences fell within a successfully genotyped, gap-free, chain and, therefore, eligible for *Pv3Rs* classification were included.

### Inclusion and Ethics

All samples in this study were derived from blood samples obtained from patients positive for malaria and collected with informed consent from the patient, or patient parent / legal guardian where individuals were less than 18 years of age. At each location, sample collection was approved by the appropriate local institutional ethics committees. The following committees gave ethical approval for the partner studies; Human Research Ethics Committee of NT Department of Health and Families and Menzies School of Health Research, Darwin, Australia (HREC-2013-1991 and HREC-2019-3288); the National Research Ethics Review Committee of Ethiopia (Ref No. EPHI6.13/4362009); College of Medicine and Health Sciences Institutional Research Ethics Review Board, Arbaminch, Ethiopia (CMHS/121733883/111); Columbia University Medical Center Institutional Review Board, New York, United States (IRB-AAAQ3503).

## RESULTS

### High-quality genotyping data on P. vivax infection pairs

In total, 978 individuals in Ethiopia were enrolled in the two clinical trials (IMPROV N = 580; OPRA N = 398), of which 292 (29.9%) experienced a recurrent infection during the 9 to 12-month follow-up period. From these clinical trials, 1,225 parasite samples were successfully genotyped from 786 participants. This included 604 samples from 444 participants enrolled in IMPROV and 621 samples across 342 participants in OPRA. Of these 786 individuals, 190 had at least one consecutive pair of genotyped samples (i.e. enrolment and recurrence 1 or consecutive recurrences) (IMPROV = 64, OPRA = 126) (**Table 1**). A total of 79 individuals experienced 2 or more genotyped recurrences (IMPROV = 16; OPRA = 63).

**Table 1:** Overview of cohort enrolment and sample genotyping.

|  | **IMPROV** | **OPRA** | **Total** |
|---|---|---|---|
| Participants enrolled in clinical trial | 580 | 398 | 978 |
| Participants with ≥1 genotyped recurrence | 444 | 342 | 786 |
| Participants with ≥1 genotyped consecutive pair | 64 | 126 | 190 |
| Participants with ≥2 genotyped recurrences | 16 | 63 | 79 |
| **Samples successfully genotyped** | **604** | **621** | **1,225** |

**Supplementary Figure S1:**
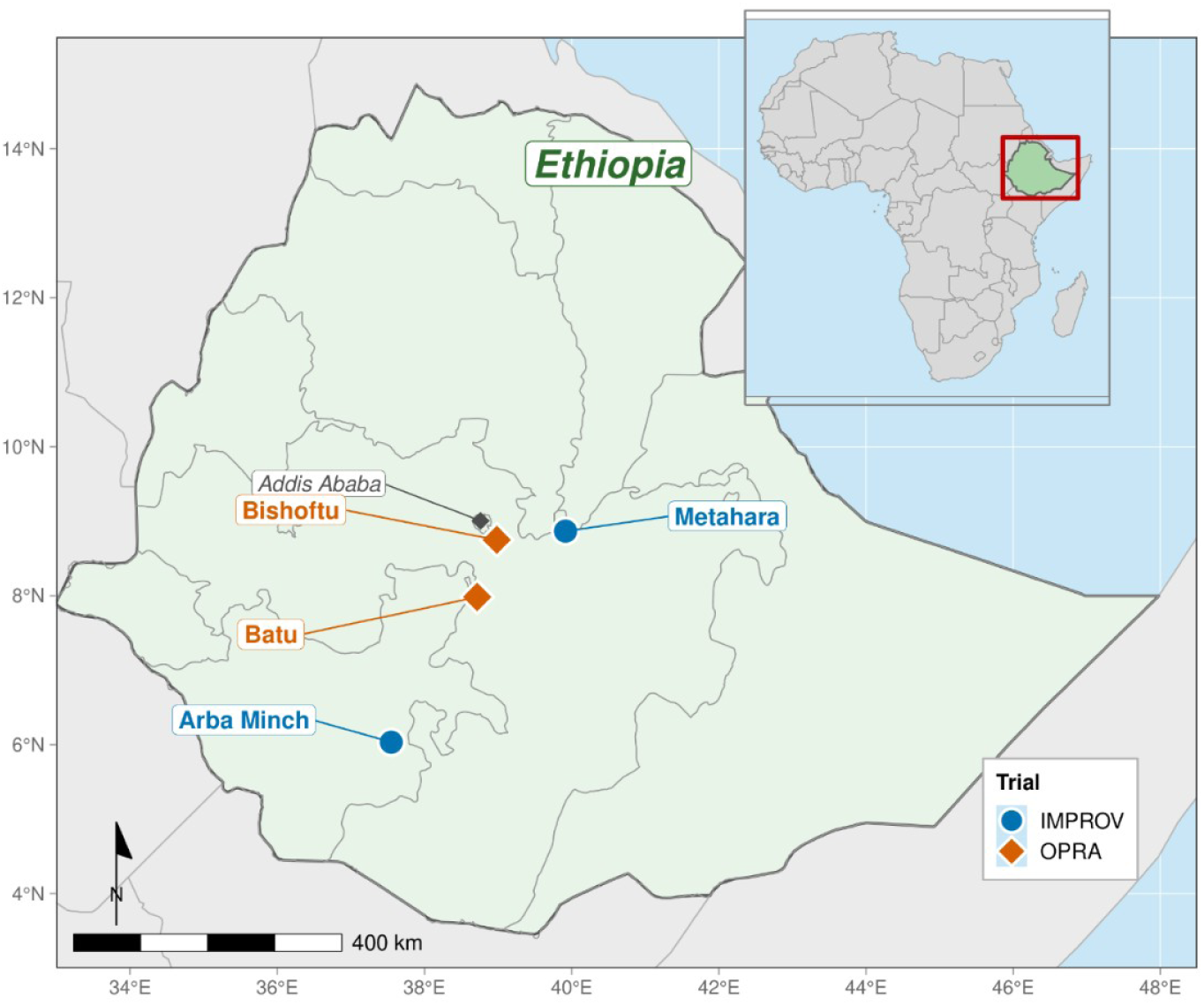
Map of IMPROV and OPRA sites

### High baseline genetic diversity with limited spatial structuring across Ethiopian P. vivax populations

Parasite populations at enrolment (Day 0) in both trials showed high genetic diversity and limited population structure, consistent with substantial gene flow across the study regions and other high transmission settings. Infection complexity was higher in IMPROV than in OPRA, with 49% (212/433) of Day 0 samples from IMPROV estimated to be polyclonal (effective multiplicity of infection [eMOI] > 1.1), compared to 25.9% (85/328) of Day 0 samples from OPRA (**Supplementary Figure S2 & S3**). The mean eMOI in IMPROV was 1.58 compared to 1.17 in OPRA. Background relatedness among Day 0 samples was low across all sites in both trials (mean pairwise IBD range = 2.9 to 4.9%), supporting the reliability of relatedness-based recurrence classification in this setting (**Supplementary Figure S3**). Polyclonal infections (eMOI > 1.1) modestly inflated these background estimates relative to monoclonal-only comparisons **(Supplementary Figure S4)**. Full site-level characterisation of allelic diversity, eMOI, background IBD, and geographic structure is provided in **Supplementary Results S1**.

IBD pairwise relatedness of within-host parasite relatedness was significantly lower in patients treated with PQ administration in both trials, consistent with the probabilistic classification results that follow (**Supplementary Figure S5**). The impact of treatment arm on eMOI per infection episode was assessed to determine if the implementation of radical cure reduced parasite diversity of the hypnozoite reservoir over time but our data did not support this (**Supplementary Figure S6**). The relatedness estimates, including stratification by day of follow-up, for all treatment arms are provided in **Supplementary Results S2**.

### Relapses driving primary recurrent infections in the absence of primaquine

Among first recurrences, 65.6% (21/32) of control arm pairs in IMPROV were classified as probable relapses, 15.6% (5/32) classified as reinfections, and 18.8% (6/32) as indeterminate (**Table 2**). There were significantly fewer classified probable relapses in the PQ arms at 12.9% (4/31), compared to 80.6% (25/31) classified as reinfection and 6.5% (2/31) as indeterminate. The classification proportions differed significantly between treatment arms (χ² = 26.9, df = 2, p = 1.45×10⁻⁶). Recurrent episodes were classified as a relapse if the *Pv3Rs* posterior probability of relapse, p(L), was ≥ 0.75 and as a reinfection if p(L) ≤ 0.25 [10,11,30]. Values between p(L) 0.25 and 0.75 were classified as indeterminate (**Figure 1**; **Supplementary Figure S7**). These thresholds require reasonably strong posterior confidence (at least 3:1 posterior odds) before assigning a definitive classification. The high posterior confidence threshold ensures that pairs with uncertain posterior support are treated as ambiguous under the model (**Supplementary Table 2**).

**Figure 1:**
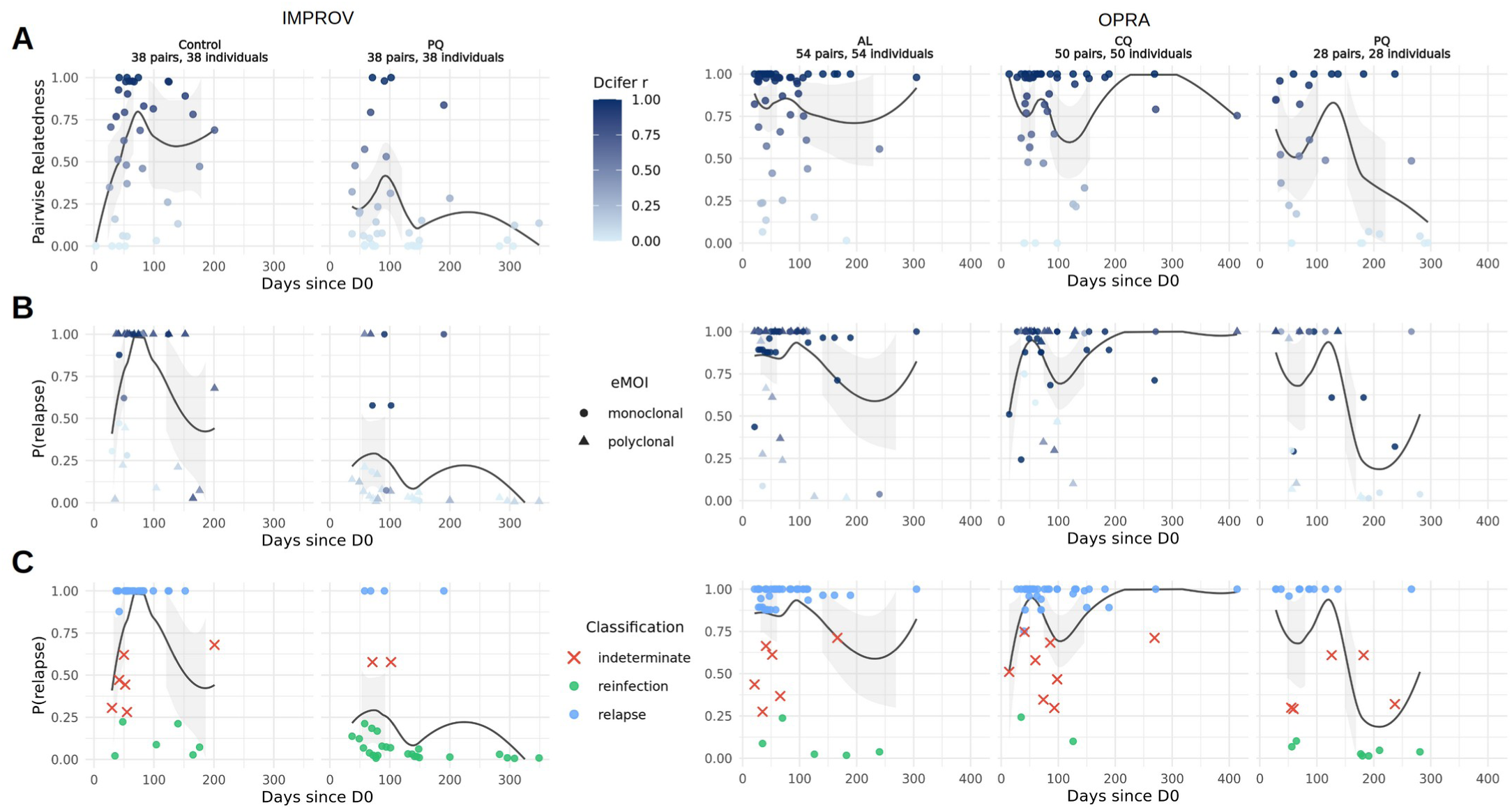
Comparison of identity-by-descent (IBD) estimates with recurrence classification probabilities over the course of IMPROV and OPRA clinical trials and follow-up period for first recurrences only (Day 0 to first recurrent infection). (A) Pairwise relatedness (IBD) estimates between Day 0 and first recurrence. (B) posterior probability of relapse, p(L). (C) Classification of first recurrences in trial treatment arms; probable relapse = p(L) ≥ 0.75, reinfection if p(L) ≤ 0.25, indeterminate when values between p(L) 0.25 and 0.75. eMOI describes clonality of recurrent infection point. Pairwise relatedness estimates represent relatedness of an isolate compared to the immediately preceding isolate within an individual.

**Supplementary figure S7:**
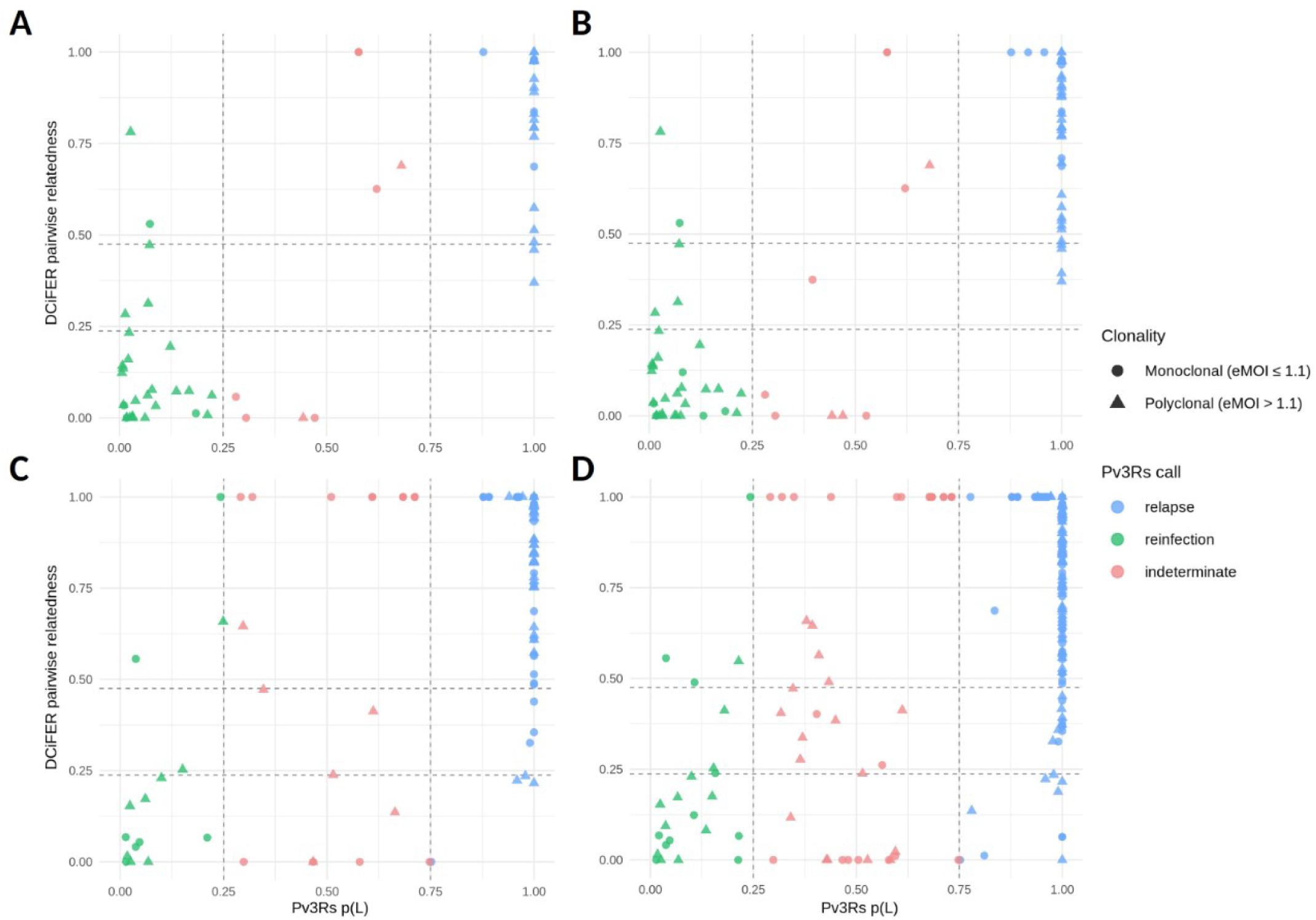
Pairwise relatedness and infection complexity compared to *Pv3Rs* informed classification using posterior probability of relapse, p(L), was ≥ 0.75 and as a reinfection if p(L) ≤ 0.25. Values between p(L) 0.25 and 0.75 classified as indeterminate. (A) First recurrences in IMPROV; (B) All recurrences in IMPROV; (C) First recurrences in OPRA; (D) All recurrences in OPRA. Dotted lines correlate to the p(L) posterior thresholds 0.25 and 0.75, and pairwise relatedness values ∼0.25 (IBD ≥ 0.2475; genetic cousins) and ∼0.50 (IBD ≥ 0.475; genetic siblings).

**Table 2:** Classification of recurrent infections from IMPROV and OPRA, grouped by treatment arm, number of recurrences, and time between Day 0 and first recurrence. Recurrent episodes classified as a relapse if the *Pv3Rs* posterior probability of relapse, p(L), was ≥ 0.75 and as a reinfection if p(L) ≤ 0.25. Values between p(L) 0.25 and 0.75 classified as indeterminate.

| First Recurrence Only |  |  |  |  |  |  |  |  |  |  |  |  |  |
| Trial | Arm | Day 0 Genotyped | Did not recur* |  | Classifiable Pairs | Relapse |  | Reinfection |  | Indeterminate |  | Mean days | Med. days |
|  |  |  | N | % |  | N | % | N | % | N | % |  |  |
|---|---|---|---|---|---|---|---|---|---|---|---|---|---|
| **OPRA** | AL | 86 | 32 | 37.2 | 54 | 43 | 79.6 | 5 | 9.3 | 6 | 11.1 | 75.8 | 56 |
|  | CQ | 95 | 40 | 42.1 | 49 | 38 | 77.6 | 2 | 4.1 | 9 | 18.4 | 92.2 | 67 |
|  | PQ arms | 147 | 122 | 83 | 24 | 12 | 50 | 7 | 29.2 | 5 | 20.8 | 120.3 | 91 |
| **IMPROV** | Control | 75 | 32 | 42.7 | 32 | 21 | 65.6 | 5 | 15.6 | 6 | 18.8 | 73.8 | 56 |
|  | PQ arms | 286 | 231 | 80.8 | 31 | 4 | 12.9 | 25 | 80.6 | 2 | 6.5 | 120.3 | 91 |
| **All Consecutive Recurrences** |  |  |  |  |  |  |  |  |  |  |  |  |  |
| **Trial** | **Arm** | **Day 0 Genotyped** | **Did not recur\*** |  | **Classified Pairs** | **Relapse** |  | **Reinfection** |  | **Indeterminate** |  |  |  |
|  |  |  | **N** | **%** |  | **N** | **%** | **N** | **%** | **N** | **%** |  |  |
| **OPRA** | AL | 86 | 32 | 37.2 | 119 | 90 | 75.6 | 10 | 8.4 | 19 | 16.0 |  |  |
|  | CQ | 95 | 40 | 42.1 | 104 | 78 | 75.0 | 4 | 3.8 | 22 | 21.2 |  |  |
|  | PQ arms | 147 | 122 | 83.0 | 40 | 25 | 62.5 | 10 | 25 | 5 | 12.5 |  |  |
| **IMPROV** | Control | 75 | 32 | 42.7 | 55 | 39 | 70.9 | 8 | 14.5 | 8 | 14.5 |  |  |
|  | PQ arms | 286 | 231 | 80.8 | 33 | 5 | 15.2 | 26 | 78.8 | 2 | 6.1 |  |  |
| **Monoclonal Consecutive Recurrences (eMOI ≤ 1.1)** |  |  |  |  |  |  |  |  |  |  |  |  |  |
| **Trial** | **Arm** | **Day 0 Genotyped** | **Did not recur\*** |  | **Classified Pairs** | **Relapse** |  | **Reinfection** |  | **Indeterminate** |  |  |  |
|  |  |  | **N** | **%** |  | **N** | **%** | **N** | **%** | **N** | **%** |  |  |
| **OPRA** | AL | 63 | 26 | 41.3 | 69 | 56 | 81.2 | 4 | 5.8 | 9 | 13 |  |  |
|  | CQ | 68 | 33 | 48.5 | 57 | 43 | 75.4 | 2 | 3.5 | 12 | 21.1 |  |  |
|  | PQ arms | 112 | 95 | 84.8 | 27 | 17 | 63.0 | 5 | 18.5 | 5 | 18.5 |  |  |
| **IMPROV** | Control | 39 | 19 | 48.7 | 15 | 9 | 60.0 | 1 | 6.7 | 5 | 33.3 |  |  |
|  | PQ arms | 144 | 120 | 83.3 | 9 | 2 | 22.2 | 5 | 55.6 | 2 | 22.2 |  |  |

In IMPROV, 57.3% (43/75) of participants in the control arm had a clinical recurrence of *P. vivax* during the 12-month follow-up period, compared to 19.2% (55/286) of participants in the treated with PQ (**Table 2**; **Supplementary Table S2**). The median number of days between the initial infection (Day 0) and first recurrence was 56 days (range 30-201) in the control arm and 91 days (range 37-349) in the PQ arms. Among first recurrences, 65.6% (21/32) of control arm pairs were classified as probable relapses, 15.6% (5/32) classified as reinfections, and 18.8% (6/32) as indeterminate. There were significantly fewer classified probable relapses in the PQ arms at 12.9% (4/31), compared to 80.6% (25/31) classified as reinfection and 6.5% (2/31) as indeterminate. The classification proportions differed significantly between treatment arms (χ² = 26.9, df = 2, p = 1.45×10⁻⁶).

In OPRA, the majority of first recurrences in the non-PQ arms (AL and CQ) were also classified as probable relapse, where 79.6% (43/54) of AL pairs and 77.6% (38/49) of CQ pairs had an estimated p(L) ≥ 0.75 (**Table 2**; **Supplementary Table S2**). In the AL arm, 9.3% (5/54) were classified as probable reinfections and 11.1% (6/54) were indeterminate (median 56 days to first recurrence). The corresponding proportions in patients in the CQ arm, were 4.1% (2/49) probable reinfections and 18.4% (9/49) indeterminate (median 67 days). In the PQ arms (ALPQ and CQPQ), the proportions of probable relapses decreased to 50% (12/24), with reinfections accounting for 29.2% (7/24) of recurrences and 20.8% (5/24) indeterminate. The classification proportions differed significantly across the three OPRA arms (χ^2^ = 12.9, df = 4, p = 0.0117). Overall participants treated with PQ had fewer clinically observed recurrences, with only 17% (25/147) of participants experiencing a recurrent infection, compared to 62.8% (54/86) and 57.9% (55/95) of participants in the AL and CQ arms, respectively.

**Supplementary Table 2:** *Pv3Rs* posterior probabilities recurrent infections from IMPROV and OPRA, grouped by treatment arm and number of recurrences.

| First Recurrence Only |  |  |  |  |  |  |  |  |  |  |  |  |
|---|---|---|---|---|---|---|---|---|---|---|---|---|
| **Trial** | **Arm** | **Day 0 Genotyped** | **Pairs** | **p(L)** |  |  | **p(I)** |  |  | **p(C)** |  |  |
|  |  |  |  | **N** | **%** | **Mean p** | **N** | **%** | **Mean p** | **N** | **%** | **Mean p** |
| **OPRA** | AL | 86 | 54 | 47 | 87.0 | 0.9409 | 7 | 13.0 | 0.8506 | 0 | 0 | 0.0296 |
|  | CQ | 95 | 49 | 43 | 87.8 | 0.9354 | 5 | 10.2 | 0.0868 | 1 | 2 | 0.0532 |
|  | PQ arms | 147 | 24 | 16 | 66.7 | 0.5981 | 8 | 33.3 | 0.3115 | 0 | 0 | 0.0904 |
| **IMPROV** | Control | 75 | 32 | 23 | 71.9 | 0.7535 | 9 | 28.1 | 0.2427 | 0 | 0 | 0.0038 |
|  | PQ arms | 286 | 31 | 6 | 19.4 | 0.2137 | 25 | 80.6 | 0.7591 | 0 | 0 | 0.0273 |
| **All Consecutive Recurrences** |  |  |  |  |  |  |  |  |  |  |  |  |
| **Trial** | **Arm** | **Day 0 Genotyped** | **Pairs** | **p(L)** |  |  | **p(I)** |  |  | **p(C)** |  |  |
|  |  |  |  | **N** | **%** | **Mean p** | **N** | **%** | **Mean p** | **N** | **%** | **Mean p** |
| **OPRA** | AL | 86 | 119 | 105 | 88.2 | 0.9328 | 14 | 11.8 | 0.8095 | 0 | 0 | 0.0337 |
|  | CQ | 95 | 104 | 84 | 84.8 | 0.8441 | 13 | 13.1 | 0.1171 | 2 | 2 | 0.0388 |
|  | PQ arms | 147 | 40 | 29 | 72.5 | 0.6882 | 11 | 27.5 | 0.2510 | 0 | 0 | 0.0608 |
| **IMPROV** | Control | 75 | 55 | 42 | 76.4 | 0.7841 | 13 | 23.6 | 0.2115 | 0 | 0 | 0.0045 |
|  | PQ arms | 286 | 33 | 7 | 21.2 | 0.2345 | 26 | 78.8 | 0.7398 | 0 | 0 | 0.0256 |
| **Monoclonal Consecutive Recurrences (eMOI ≤ 1.1)** |  |  |  |  |  |  |  |  |  |  |  |  |
| **Trial** | **Arm** | **Day 0 Genotyped** | **Pairs** | **p(L)** |  |  | **p(I)** |  |  | **p(C)** |  |  |
|  |  |  |  | **N** | **%** | **Mean p** | **N** | **%** | **Mean p** | **N** | **%** | **Mean p** |
| **OPRA** | AL | 63 | 69 | 65 | 94.2 | 0.8884 | 4 | 5.8 | 0.0548 | 0 | 0 | 0.0568 |
|  | CQ | 68 | 57 | 51 | 89.5 | 0.8506 | 4 | 7 | 0.0835 | 2 | 3.5 | 0.0659 |
|  | PQ arms | 112 | 27 | 21 | 77.8 | 0.7072 | 6 | 22.2 | 0.2028 | 0 | 0 | 0.0901 |
| **IMPROV** | Control | 39 | 15 | 11 | 73.3 | 0.7344 | 4 | 26.7 | 0.2493 | 0 | 0 | 0.0164 |
|  | PQ arms | 144 | 9 | 4 | 44.4 | 0.3913 | 5 | 55.6 | 0.5147 | 0 | 0 | 0.0940 |

### Relapse is the dominant mechanism of recurrence in the absence of radical cure

When considering all consecutive recurrence pairs within an individual, 70.9% (39/55) of recurrences in participants enrolled in IMPROV, not treated with PQ, were classified as probable relapses, compared to 15.2% (5/33) of participants treated with PQ (**Table 2**; **Figure 3**). Probable reinfections [p(L) ≤ 0.25] accounted for 14.5% (8/55) of recurrences in the control arm and 78.8% (26/33) in the PQ arms (χ² = 36.2, df = 2, p = 1.40×10⁻⁸). In OPRA, 75.6% (90/119) of AL pairs and 75.0% (78/104) of CQ pairs were classified as probable relapses, compared to 62.5% (25/40) of PQ arm pairs (χ² = 19.1, df = 4, p = 0.0007). Relapse estimates in the PQ arms of OPRA (62.5%; 25/40) were higher than that in the PQ arms of IMPROV (15.2%; 5/33). This may reflect the lower dose of primaquine administered in OPRA or reduced adherence to the full PQ regimen, as only the initial treatment was supervised at the time of enrolment. In the OPRA trial, the proportion of participants with probable relapses rose from 50.0% (12/24), when only considering first recurrences, to 62.5% (25/40) when accounting for all recurrences.

**Figure 2:**
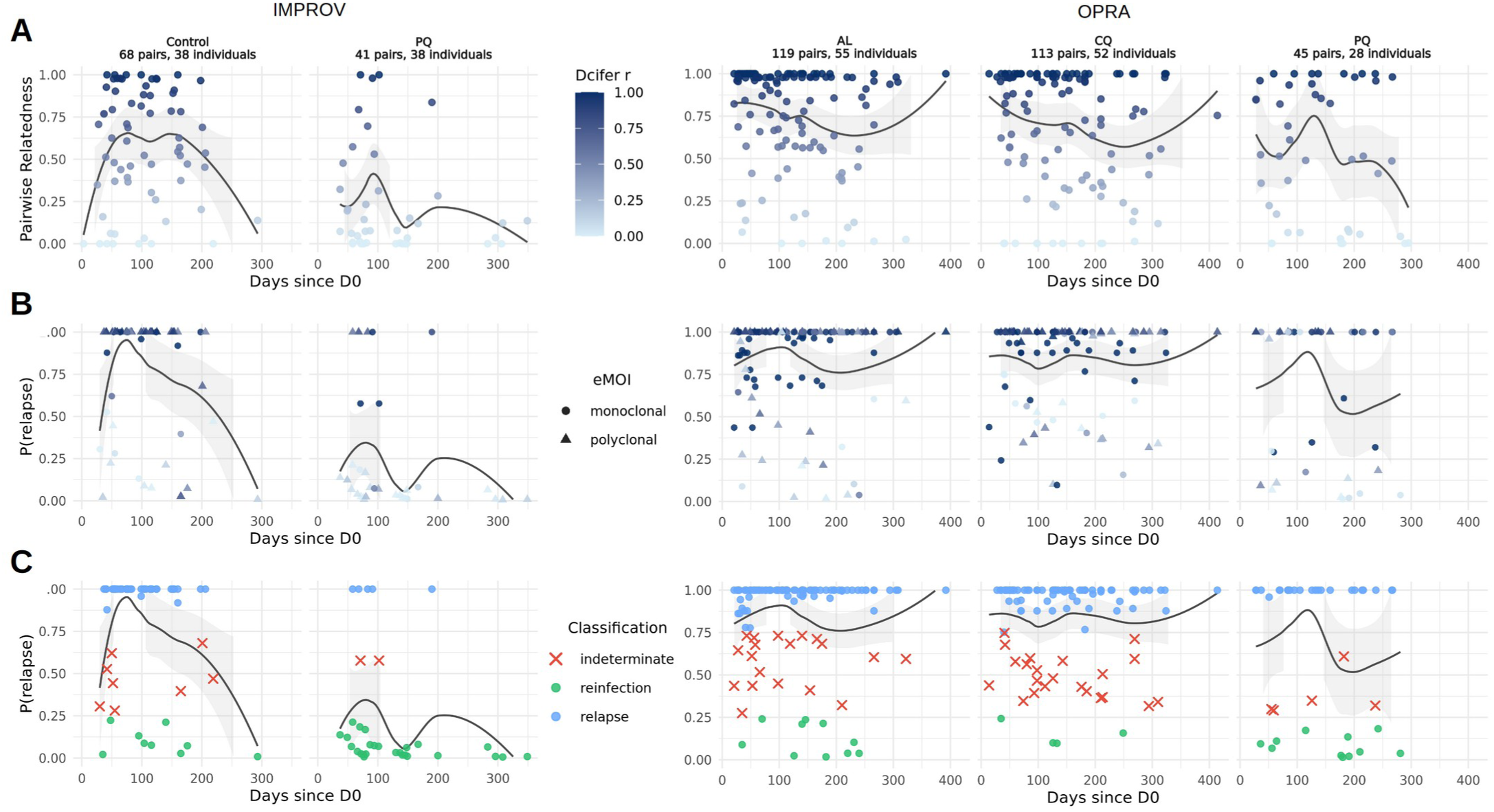
Comparison of identity-by-descent (IBD) estimates with recurrence classification probabilities over the course of IMPROV and OPRA clinical trials and follow-up period for all recurrences. (A) Pairwise relatedness (IBD) estimates between Day 0 and all recurrences. (B) *Pv3Rs* posterior probability of relapse, p(L). (C) Classification of first recurrences in trial treatment arms; probable relapse if p(L) ≥ 0.75, reinfection if p(L) ≤ 0.25, indeterminate when values between p(L) 0.25 and 0.75. eMOI describes clonality of recurrent infection, while pairwise relatedness estimates represent relatedness of an isolate compared to the immediately preceding isolate within an individual.

**Figure 3:**
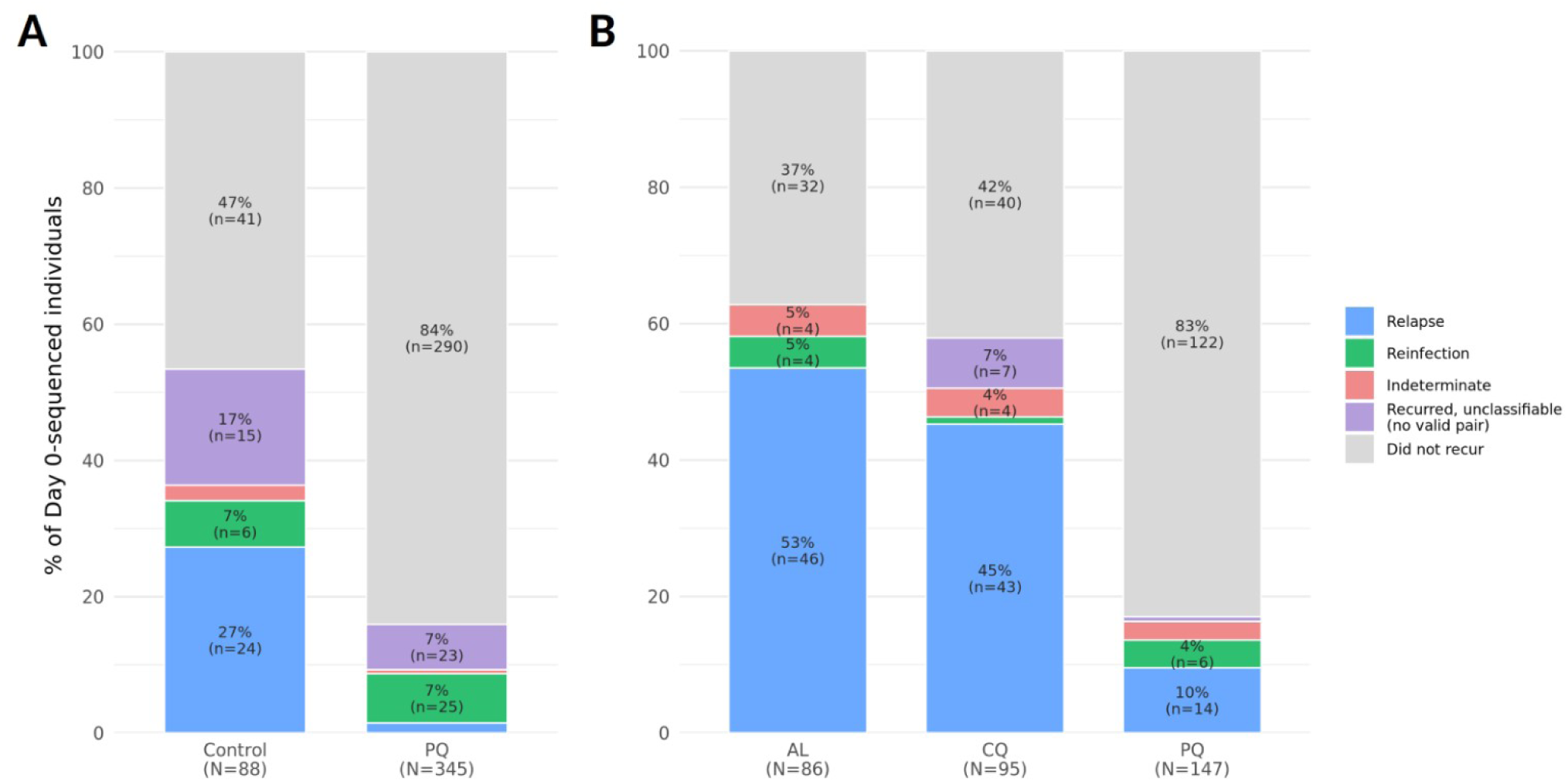
Individual outcomes informed by classified recurrences from (A) IMPROV and (B) OPRA grouped by treatment arm. Relapse if the *Pv3Rs* p(L) ≥ 0.75; reinfection p(L) ≤ 0.25; Indeterminate, values between p(L) 0.25 and 0.75. Individuals that recurred but were missing genotyping from a recurrence were labelled unclassifiable.

In OPRA participants not treated with PQ, 74.0% (37/50) of individuals with polyclonal infection at enrolment had recurrence, compared to 55.0% (72/131) of patients with monoclonal infection at enrolment (Odds Ratio [OR] = 2.35; 95% CI 1.14 – 4.83; p = 0.020) (**Supplementary Table S3 and Figure S8**). In IMPROV, the trend was similar (60.0% (24/40) and 47.9% (23/48) respectively), but this did not reach statistical significance (OR = 1.63; 95% CI 0.70 - 3.81; p = 0.26).

When considering only probable relapses (*Pv3Rs* p(L) ≥ 0.75), OPRA showed a modest significant association between polyclonality at enrolment and a higher count of probable relapse episodes (ratio = 1.39; 95% CI 1.01 – 1.92; p = 0.044). This was not replicated in IMPROV (ratio = 1.14; 95% CI 0.61 – 2.14, p = 0.68). No significant association was observed when pooling all polyclonal individuals at the time of enrolment in the non-PQ arms of IMPROV and OPRA, with 42.4% (72/170) of monoclonal Day 0 individuals experiencing a probable relapse versus 47.7% (41/86) of polyclonal Day 0 individuals (count ratio = 1.20; 95% CI 0.81 – 1.76; p = 0.358).

In IMPROV, 72.7% (64/88) of participants with recurrences had at least one isolate which was polyclonal, whereas in OPRA, the majority of recurrence pairs were monoclonal (59.8%; 153/256) (**Supplementary Table S3; Supplementary Figure S8**). To assess the robustness of *Pv3Rs* when classifying polyclonal infections, given the impact of polyclonality on relatedness, analyses were repeated following restriction to only monoclonal pairs (eMOI ≤ 1.1). Classification proportions in OPRA were broadly consistent between monoclonal-only and polyclonal-involving pairs (χ² = 1.50, df = 2, p = 0.472) (Table X). In IMPROV, monoclonal-only sample sizes were too small to be statistically meaningful; results are presented in **Supplementary Results S3.**

**Supplementary Table 3:** Impact of polyclonality at time of enrolment (Day 0), using eMOI estimates, on likelihood of recurrence in IMPROV and OPRA. Monoclonal (eMOI ≤ 1.1); Polyclonal (eMOI > 1.1).

| **Trial** | **Arm** | **Monoclonal D0 Recur** |  | **Polyclonal D0 Recur** |  | **Monoclonal D0 ≥1 Relapse\*** |  | **Polyclonal D0 ≥1 Relapse** |  |
|---|---|---|---|---|---|---|---|---|---|
|  |  | **%** | **N** | **%** | **N** | **%** | **N** | **%** | **N** |
| **IMPROV** | Control (CQ) | 47.9% | 23/48 | 60.0% | 24/40 | 30.8% | 12/39 | 33.3% | 12/36 |
| **OPRA** | AL | 58.7% | 37/63 | 73.9% | 17/23 | 50.8% | 32/63 | 60.9% | 14/23 |
| **OPRA** | CQ | 51.5% | 35/68 | 74.1% | 20/27 | 41.2% | 28/68 | 55.6% | 15/27 |
| **Combined** | **Non-PQ** | **53.1%** | **95/179** | **67.8%** | **61/90** | **42.4%** | **72/170** | **47.7%** | **41/86** |

\*Relapse defined as a posterior probability of relapse, p(L) ≥ 0.75.

**Supplementary Figure S8:**
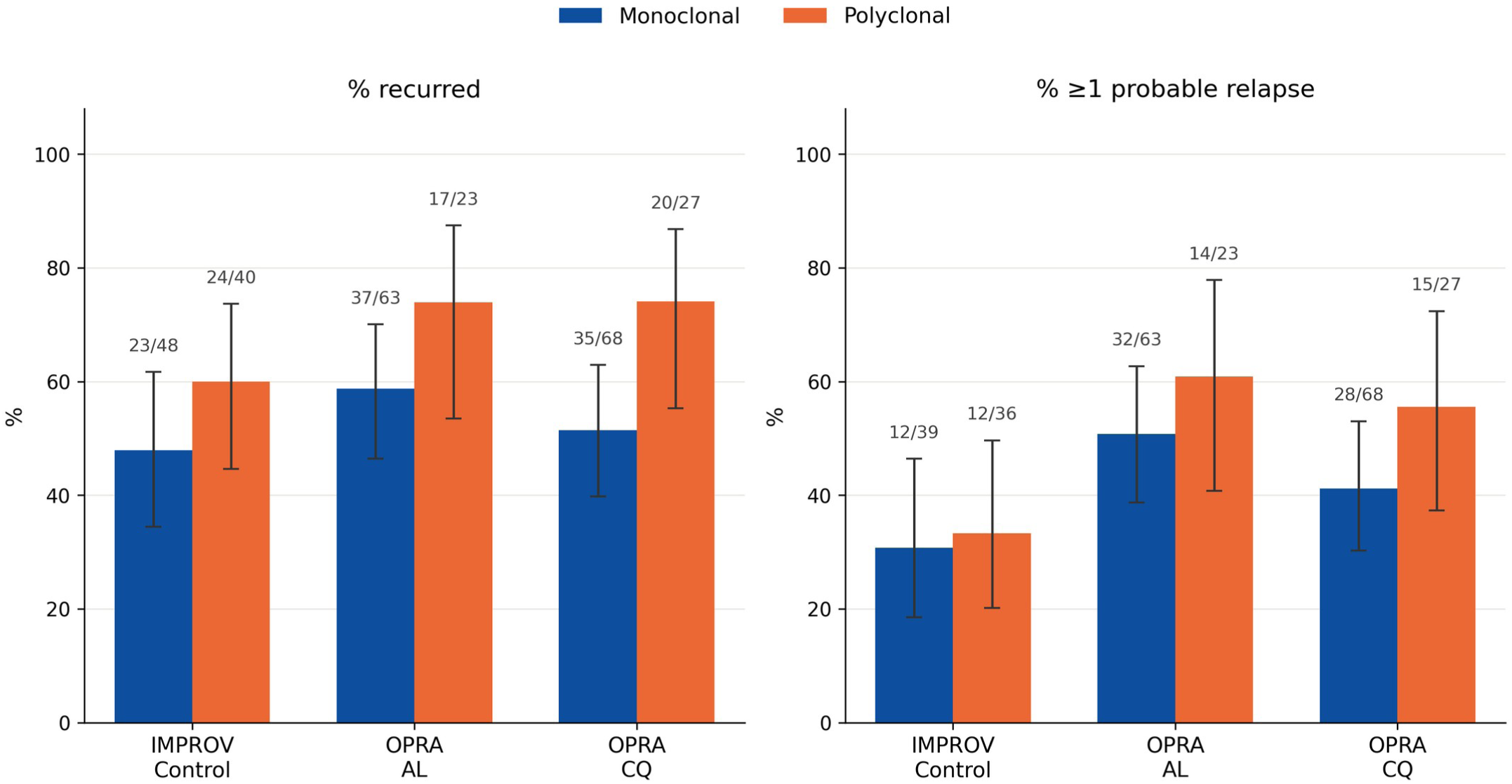
Impact of polyclonality at time of enrolment (Day 0) on likelihood of recurrence in IMPROV and OPRA. Monoclonal (eMOI ≤ 1.1); Polyclonal (eMOI > 1.1); 95% confidence interval (Wilson method).

### The individual burden of persistent relapse infections

In IMPROV, only 19.2% (55/286) of genotyped participants treated with PQ, who could be genotyped, had a recurrent infection, compared to 57.3% (43/75) of those in the control arm. Only 1.7% (5/286) of PQ arm participants had classifiable probable relapses, with the majority of recurrences attributed to probable reinfections (**Figure 3**, **Supplementary Figure S9**). The time between enrolment and first recurrence was also longer in the PQ arms (median 91 days) than in the control arm (median 56 days). This pattern is consistent with successful clearance of the hypnozoite reservoir resulting in residual recurrences arising predominantly from new infections rather than treatment failure. The maximum number of relapses within control arm participants was 4 over a period of 206 days (mean = 1.22; median = 1; N = 32), compared to only 1, over 190 days, in the PQ arms (mean = 0.16; median = 0; N = 32). A significant difference in per-individual relapse counts was observed between the control arm and the PQ arms (H = 74.4, p = 6.3×10⁻¹⁸).

In OPRA, a similar pattern was observed, with only 17.0% (25/147) of participants treated with PQ experiencing a recurrence, compared to 62.8% (54/86) of those treated with AL and 57.9% (55/95) of those treated with CQ. Of the 147 participants treated with PQ, the majority of recurrences were attributed to probable reinfection with only 9.5% (14/147) classified as probable relapse. The time between enrolment and first recurrence was also longer in participants treated with PQ (median 91 days) than in those treated with either AL (median 56 days) or CQ arms (median 67 days). The findings are similar to that in the IMPROV trial, although the effect was less pronounced in OPRA, likely reflecting OPRA’s lower-dose and partially unsupervised primaquine regimen (see Discussion). The maximum number of relapses within an individual was 6 over 392 days in the AL arm (mean = 1.69; median = 1; N = 54), 5 over 295 days in the CQ arm (mean = 1.54; median = 1; N = 48), and 3 over 201 days in the PQ arms (mean = 1.04; median = 1; N = 24); (H = 58.4, p = 2.1×10⁻¹³).

**Supplementary Figure S9:**
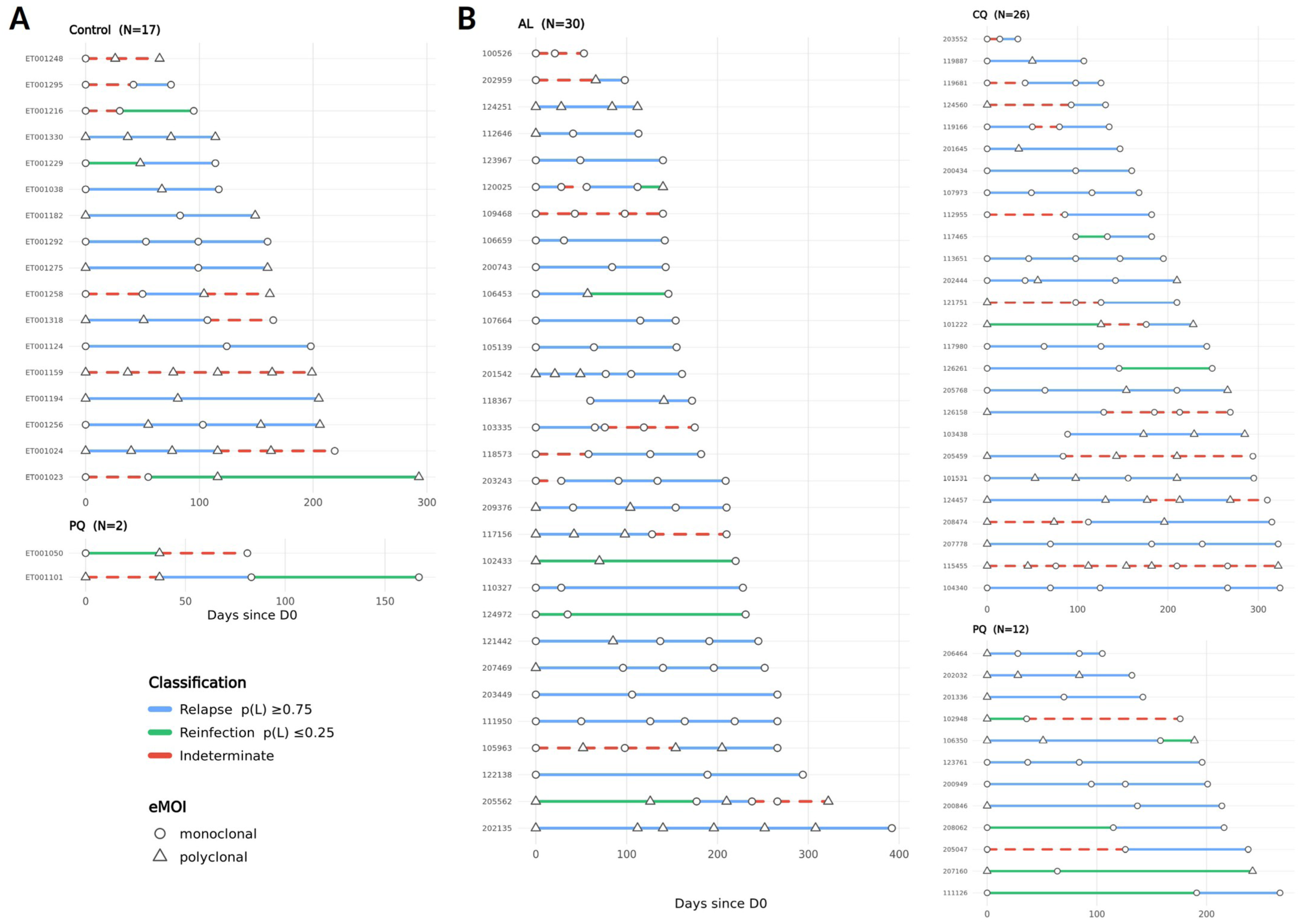
Within-individual recurrence classification timelines, grouped by trial and treatment arm; (A) IMPROV; (B) OPRA. Minimum of 2 consecutive genotyped recurrences (e.g. Day 0, Rec 1, Rec 2, etc.).

## DISCUSSION

Recurrent *P. vivax* malaria remains one of the greatest barriers to achieving malaria elimination in Asia, the Americas, and the Horn of Africa. Disentangling relapse from reinfection has undermined our ability to quantify the impact of relapses on the burden of the parasite and its onward transmission [3,31]. Using microhaplotype sequencing and a probabilistic classification model (*Pv3Rs*) to assess relatedness relationships within participants enrolled in two clinical trials, we provide molecular evidence that, in the absence of radical cure, relapse, not reinfection, is the dominant driver of recurrent *P. vivax* in Ethiopia [11,13,19,30]. The burden of relapse in Ethiopia likely carries a substantial, clinical cost at both the individual and community level [32].

The trial sites included in our analysis spanned geographic regions and transmission periods. The OPRA trial was conducted in central Ethiopia between 2012-2014, a period of moderate transmission, while the IMPROV study was conducted in 2017 to 2018, during a period of low incidence of malaria nationally. The southern area of Arba Minch has, however, historically been associated with higher transmission intensity [1,18,29]. Our analysis characterised the baseline genetic epidemiology of parasite populations across the trial study sites, demonstrating high genetic diversity and limited geographically linked population structure, consistent with regions that exhibit moderate to high malaria transmission and regular gene flow. Infection complexity, or the presence of multiple genetically detectable parasite clones, was greater amongst the IMPROV sites at time of enrolment. This is supported by population-level reinfection rates in the control arms which were estimated to be 9.3% (7/75) in IMPROV compared to 5.5% (10/181) in OPRA. The genetically homogeneous populations, with polyclonal infections inflating background relatedness estimates, can make relapse and reinfection difficult to distinguish by descent alone. Hence the importance of explicitly accounting for infection complexity within any relapse classification framework applied to high-transmission settings.

The reduction in probable relapses between the control and PQ arms in both IMPROV and OPRA mirrored the trials’ published incidence risks [19,21]. This concordance between independently derived clinical and molecular estimates demonstrates consistency in the classification approach and supports the interpretation that clinically observed reductions in recurrence during these trials was primarily driven by hypnozoite clearance. In both trials, the majority of recurrences in the control arms were classified as probable relapse, rather than reinfection, with this proportion increasing when considering multiple recurrence episodes within a single individual, particularly in the PQ arms. This is consistent with a growing body of evidence that relapse may be the primary cause of recurrence even in a number of settings with varying levels of transmission intensity [33–35]. In a cohort study conducted between 2021 and 2024, Cambodian participants were treated for *P. vivax* malaria and then relocated from a malaria endemic area to a transmission-free zone; the majority of individuals (81%) relapsed within 90 days, often repeatedly [34]. In the same study, even low-density, asymptomatic, relapses were able to infect mosquitoes in laboratory feeding assays. Although we could not directly assess transmission potential in our analysis, the evidence from Cambodia suggests that the persistent relapses documented in Ethiopia, particularly those that may have gone undetected outside a clinical trial setting, will likely sustain transmission in a community.

Within the same Cambodian cohort, individuals with polyclonal infections at the point of enrolment had a higher relapse count over the follow-up period [34]. Our findings in OPRA mirror this, with individuals polyclonal at enrolment exhibiting significantly greater odds of subsequent recurrence, and a higher count of probable relapse, than monoclonal individuals; this was independent of whether individuals received AL or CQ. A similar trend was observed in IMPROV but did not reach statistical significance. Our findings lend some support to the interpretation that infection complexity at presentation may reflect of local transmission intensity and be a proxy for size and diversity of an individual’s hypnozoite reservoir and risk of subsequent relapse. However, this interpretation should be treated with caution since polyclonal infections were themselves shown to inflate background relatedness estimates. Fully resolving genetic identity within polyclonal recurrences would require haplotype-deconvolved whole-genome data, which was beyond the scope of this study but represents a valuable topic for future research.

Our findings have wider implications on the burden of untreated relapses at the individual-level. Many individuals experienced persistent probable relapses spanning months, corresponding to clinical episodes which have a significant cumulative impact on wellbeing, long-term health outcomes, and productivity [3,19,21,36]. Conservative radical cure policy in Ethiopia has been shaped by the limited availability of G6PD testing at the point-of-care, and the resulting safety constraints given the risk of haemolysis in G6PD-deficient individuals [37,38]. The availability of novel point of care diagnostics for G6PD deficiency have potential to impact policy by identifying patients who can safely tolerate tafenoquine and higher dose primaquine regimens, however this comes at an increased cost to health systems [32]. Our results are not intended to minimise safety concerns, but to provide previously unavailable molecular evidence that not only highlight the true burden of the untreated hypnozoite reservoir in Ethiopia. but also add weight to an economic analysis that will inform policy and cost allocation decisions for the expansion of safe radical cure in Ethiopia and other *P. vivax* endemic areas [18].

Our analysis has a number of limitations. Both the OPRA and IMPROV datasets are subject to sampling bias as it was not possible to genotype every recurrence reported during the follow-up period. Furthermore, individual adherence to a complete course of PQ was only semi-supervised for the first recurrence in OPRA, and not for subsequent recurrences, where in IMPROV all treatments were supervised [19,21,23]. Poor adherence of unsupervised 14-day PQ regimens is likely to attenuate treatment efficacy in OPRA patients with recurrent episodes of *P. vivax*. When interpreting recurrence classifications, the conservative reinfection prior used in the *Pv3Rs* model (λ = 0.002/day) puts our estimated probable relapse proportions as lower-bound estimates [29]. The model assigns an increasingly higher prior probability of reinfection as the interval since the preceding infection grows (i.e. passage of time). This is due to the higher suggested probability that an infection should more likely be the result of reinfection as time passes.

The impact of polyclonality on classification could not be assessed robustly in the IMPROV trial given the limited number of monoclonal pairs. However, classification proportions in OPRA were consistent between monoclonal and polyclonal pairs. The large proportion of polyclonal pairs within IMPROV also resulted in more computational failures when using *Pv3Rs* to estimate probable relapses. This occurred despite accounting for pairs exceeding the explicit *Pv3Rs* computational limit (i.e. multiplicity of infection exceeding 8 between any pairs) using a stepwise comparison fallback [11].

In a small number of pairs that were genetically identical, the posterior probability of relapse fell within the indeterminate range. In these cases, nearly all posterior mass was distributed between relapse and recrudescence, rather than reinfection, reflecting competition between these two explanations for genetically identical parasites at short inter-episode intervals. This was not due to uncertainty in the underlying pairwise genetic relatedness.

A handful of later recurrences showed no genetic relatedness to the immediately preceding episode but met the threshold to be classified as probable relapses. We interpret this as evidence that some individuals harbour more than one genetically distinct hypnozoite lineage simultaneously, acquired from separate mosquito bites. When a different lineage activates than the one responsible for the previous episode, the two samples will appear unrelated by pairwise comparison alone, despite the new episode likely arising from that individual’s own parasite reservoir rather than a new infection. *Pv3Rs* uses the full recurrence timeline and background reinfection rate to make this distinction. Unexpectedly low pairwise relatedness in probable relapses could also reflect half-sibling misspecification under the *Pv3Rs*’ two-parent assumption [30]. A dedicated time-to-event model, incorporating time-sensitive covariates, would remove dependence on the rigid, fixed time-window, priors used in this study and allow more informed flexibility given the target population.

## CONCLUSIONS

Our study provides molecular evidence that, in the absence of radical cure, hypnozoite-derived relapse, not reinfection, is likely the dominant source of recurrent *P. vivax* infection in Ethiopia. The concordance between our genetic observations and the independent clinical trial efficacy data supports the validity of the approach and the interpretation that the hypnozoite reservoir may be a substantially underappreciated contributor to the burden of *P. vivax* in Ethiopia. Our findings support expanding safe, effective, radical cure implementation in the region and regions with similar population characteristics and transmission profiles.

## SUPPLEMENTARY RESULTS

### Supplementary Results S1. Baseline genetic diversity and spatial structuring across Ethiopian *P. vivax* populations

Within-host diversity was measured using the effective multiplicity of infection (eMOI), which had high concordance with naive complexity of infection (COI) across all sites (**Supplementary Figure S1**). An eMOI mean > 1.1 was used to identify infections as polyclonal. More polyclonal infections were observed amongst the enrolment samples in IMPROV, (49% [212/433], mean eMOI = 1.58), with a mean eMOI of 1.6 in Arba Minch (49.3%; 178/361) and 1.48 in Metehara (47.2%; 34/72). In OPRA (25.9% [85/328], mean eMOI = 1.17), with a mean eMOI of 1.17 in Batu (31%; 27/87) and 1.17 (24.1%; 58/241) in Bishoftu (**Supplementary Figure S2**).

Allelic diversity within Day 0 samples did not vary considerably by collection site, with a mean of 6.2 alleles per locus in Arba Minch (median distinct alleles per locus = 5; minimum = 1, maximum = 19; N = 361) and a mean of 5.8 in Metehara (median 5.0; min = 1; max = 18; N = 72) within IMPROV (Supp/Figure X). Within OPRA, the mean number of alleles per locus was 6.4 in Bishoftu (median 5.0; min = 1; max = 27; N = 241) and 5.6 in in Batu (median 5.0; min = 1; max = 23; N = 87). Within-host polyclonality, or the average number of loci per sample carrying more than one allele, was highest in Arba Minch (mean 18.8 loci >1 allele out of 93 loci; 20.2%; N = 361 samples) and Metehara (17/93 loci; 18.3%; N = 72 samples), and lowest in Batu (7.3/93 loci; 7.9%; N = 87 samples) and Bishoftu (6.5/93 loci; 7%; N = 241 samples), consistent with observed eMOI patterns (**Supplementary Figure S2**).

**Supplementary Figure S2:**
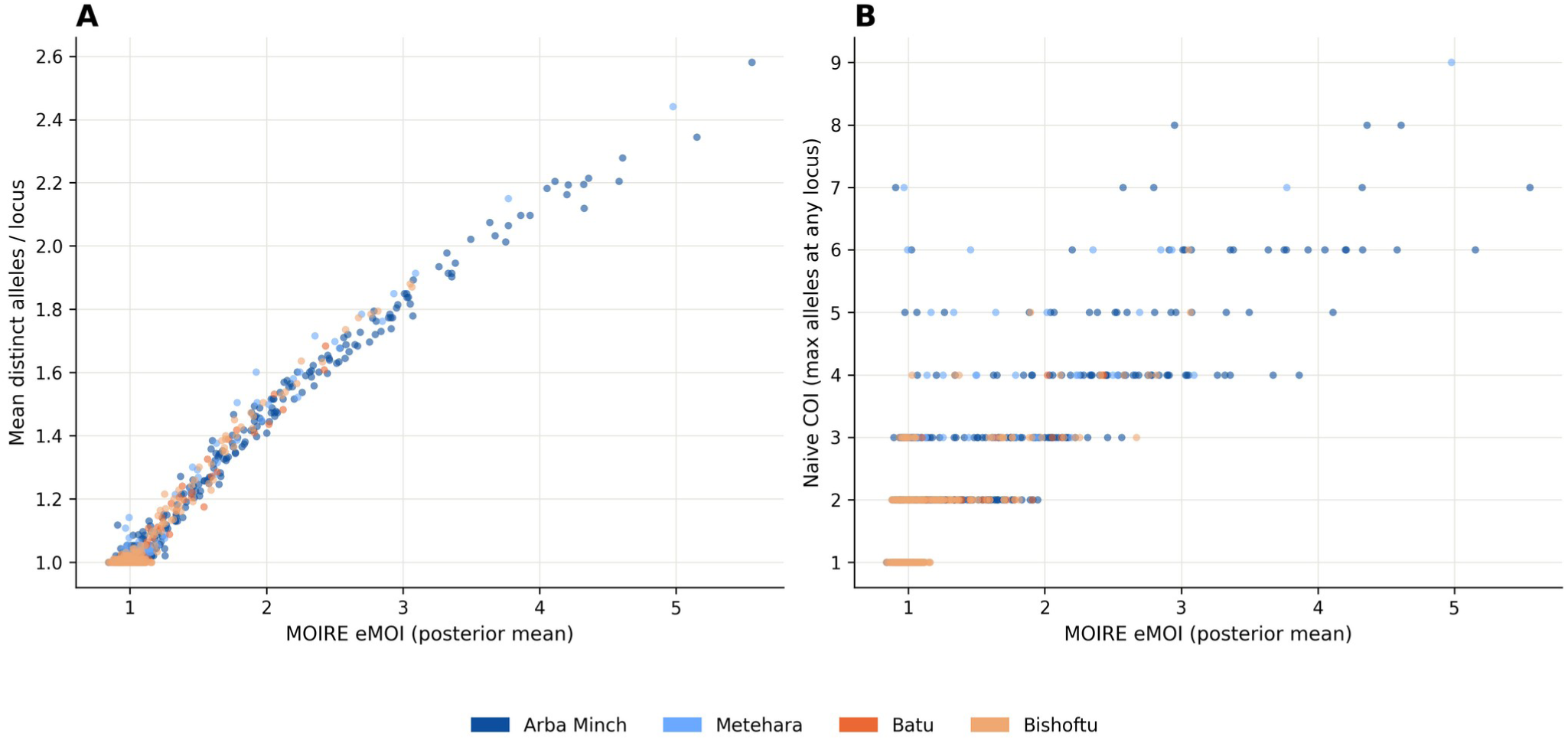
Assessing infection complexity across IMPROV and OPRA. (A) effective multiplicity of infection (eMOI) estimates compared to the mean number of alleles per locus by trial site. (B) eMOI compared to naive COI (the maximum number of alleles at any locus).

Identity-by-descent (IBD) within Day 0 samples was low within collection sites, with median relatedness of 0.002% in Arba Minch (mean 4.3%; IQR 0 – 6.7%; N = 361) and 1.8% in Metehara (mean 4.9%; IQR 0 – 7.3%; N = 72 samples) in the IMPROV trial. In OPRA, the median IBD was 0% in Batu (mean 3.1%; IQR 0 – 4.5%; N = 87 samples) and in Bishoftu (mean 2.9%; IQR 0 – 4.0%; N = 241) (**Supplementary Figure S2**). IBD values at all sites ranged from a minimum of 0% to a maximum of 100%. When restricted to monoclonal Day 0 samples (eMOI ≤ 1.1), median IBD was 0% in Arba Minch (mean 2.9%; IQR 0 – 4.3%; N = 183) and 0.3% in Metehara (mean 3.3%; IQR 0–5.2%; N = 38 samples) in IMPROV. Median IBD remained 0% in Batu (mean 2.9%; IQR 0 – 4.3%; N = 60 samples) and Bishoftu (mean 2.7%; IQR 0–3.7%; N = 183 samples). Reductions in IBD between monoclonal and polyclonal comparisons suggest polyclonality inflates background relatedness estimates.

**Supplementary Figure S3:**
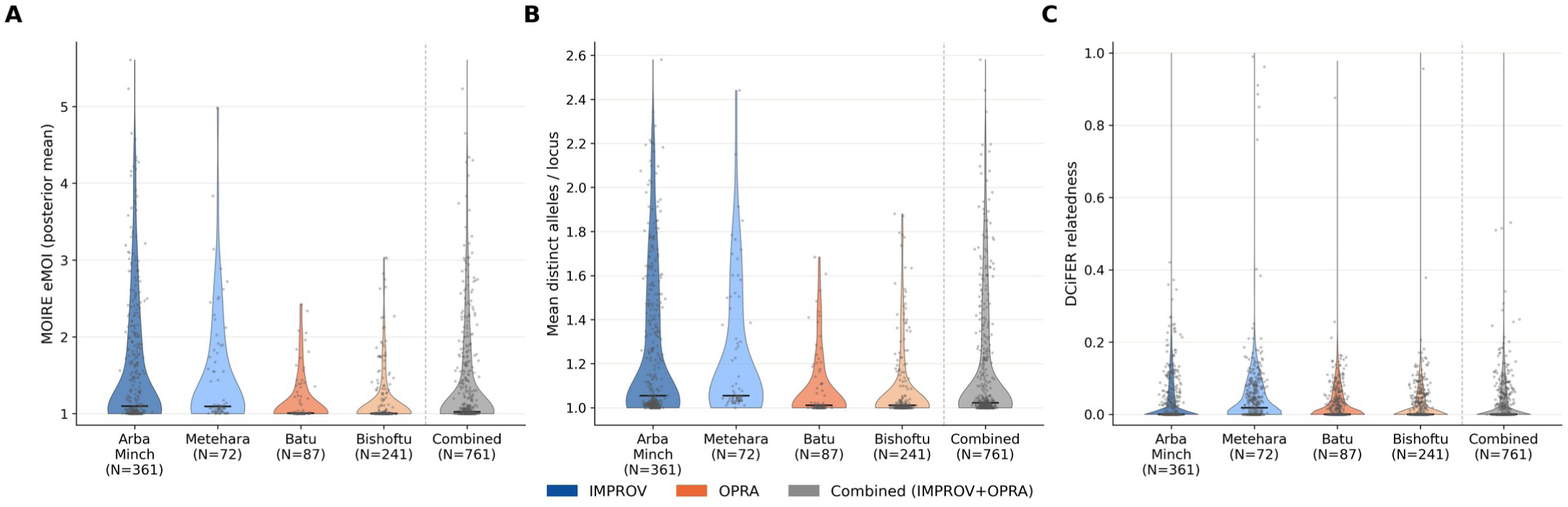
Characterisation of the baseline genetic diversity of parasite populations across IMPROV and OPRA. (A) Distribution of eMOI posterior mean. (B) Allelic diversity, the mean number of distinct alleles per locus. (C) Pairwise relatedness (identity-by-descent) per site.

**Supplementary Figure S4:**
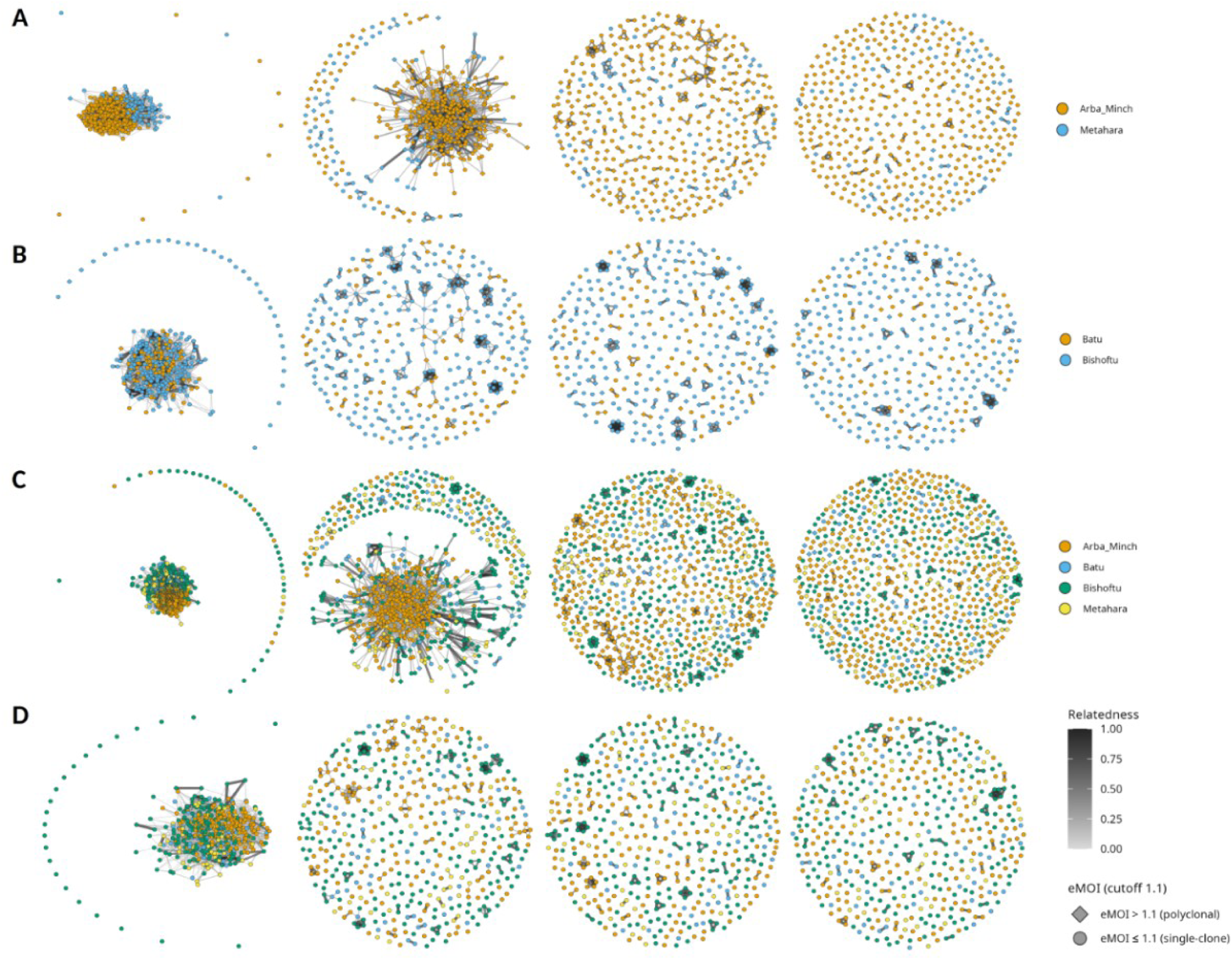
Identity-by-descent (IBD) network plot of baseline parasite populations within IMPROV and OPRA (IBD Thresholds [left to right]: 0.125, 0.25, 0.50, 0.95). (A) All Day 0 samples from IMPROV study sites: Arba Minch and Metehara. (B) All Day 0 samples from OPRA study sites: Batu and Bishoftu. (C) All Day 0 samples from both IMPROV and OPRA. (D) All monoclonal (eMOI ≤ 1.1) Day 0 samples from both IMPROV and OPRA.

Network analysis of pairwise IBD among Day 0 samples showed no clear geographic clustering by site at any relatedness threshold, consistent with gene flow across the study regions (**Supplementary Figure S3**). Geographic structure was minimal in OPRA, with same-site Day 0 pairs showing similar mean relatedness to different-site (same-site mean = 2.95%; cross-site mean = 2.63%; Mann-Whitney U p = 0.032). IMPROV same-site pairs showed somewhat higher background relatedness than cross-site pairs (same-site mean = 4.33%; cross-site mean = 2.11%; p < 2.2×10-16) but remained largely genetically unrelated. This was further supported by neighbour-joining analysis (Supp. Figures X). These results supported pooling of the sites within each trial to enhance sample size. Due to varying baseline levels of polyclonality between IMPROV sites (e.g. Arba Minch and Metehara), and the small number of infection pairs genotyped from Metehara (N = 6 pairs), the Metehara data was not grouped with data from Arba Minch and was removed from downstream analyses. Differing polyclonal estimates between the sites may skew baseline allele frequency calculations and, therefore, classification estimates.

### Supplementary Results S2. Within-host diversity and the impact of radical cure on parasite relatedness

For downstream analyses, IMPROV was restricted to only samples from Arba Minch given the small sample size of recurrent episode pairs in Metehara (N = 6 pairs). For OPRA, Batu and Bishoftu were analysed together as no meaningful site-specific differences in relatedness were observed in Day 0 samples. Within-host relatedness was assessed as the pairwise IBD between each recurrent episode and the immediately preceding infection episode, referred to as consecutive pairs (i.e. Day 0 to Recurrence 1, Recurrence 1 to Recurrence 2, and so on). This framework aligns with the biological question of whether each new recurrence is genetically continuous with its most recent predecessor.

Within-host parasite relatedness differed markedly between treatment regimens in both clinical trials (**Supplementary Figure S5**). As expected, relatedness was lower in the primaquine arms of both trials, consistent with the concept that hypnozoite clearance reduces relapse-driven recurrences. This effect was more pronounced in IMPROV, where participants received high-dose primaquine (PQ), than in OPRA, which utilised low-dose PQ, and is consistent with the differing PQ regimens between trials. Within IMPROV, the mean IBD of consecutive pairs in the control arm was 58% (N = 67; 37 individuals; interquartile range [IQR] 36.8 – 89.7%), compared to 23% in the PQ arms (N = 41; 38 individuals; IQR 0.6 – 31.3%). Pairwise Mann-Whitney U test confirmed a significant difference in relatedness within the Control arm and PQ treatment arms (p = 1.1×10⁻⁵). Restricting to monoclonal pairs (eMOI ≤ 1.1) did not produce comparable mean IBD estimates due to insufficient sample sizes (Control 62% [N = 15; 10 individuals; IQR 21.6 – 97.8%]; PQ arms 50% [N = 9; 9 individuals; IQR 3.4 – 98.0%]). This reflects the high baseline polyclonality of this population.

**Supplementary Figure S5:**
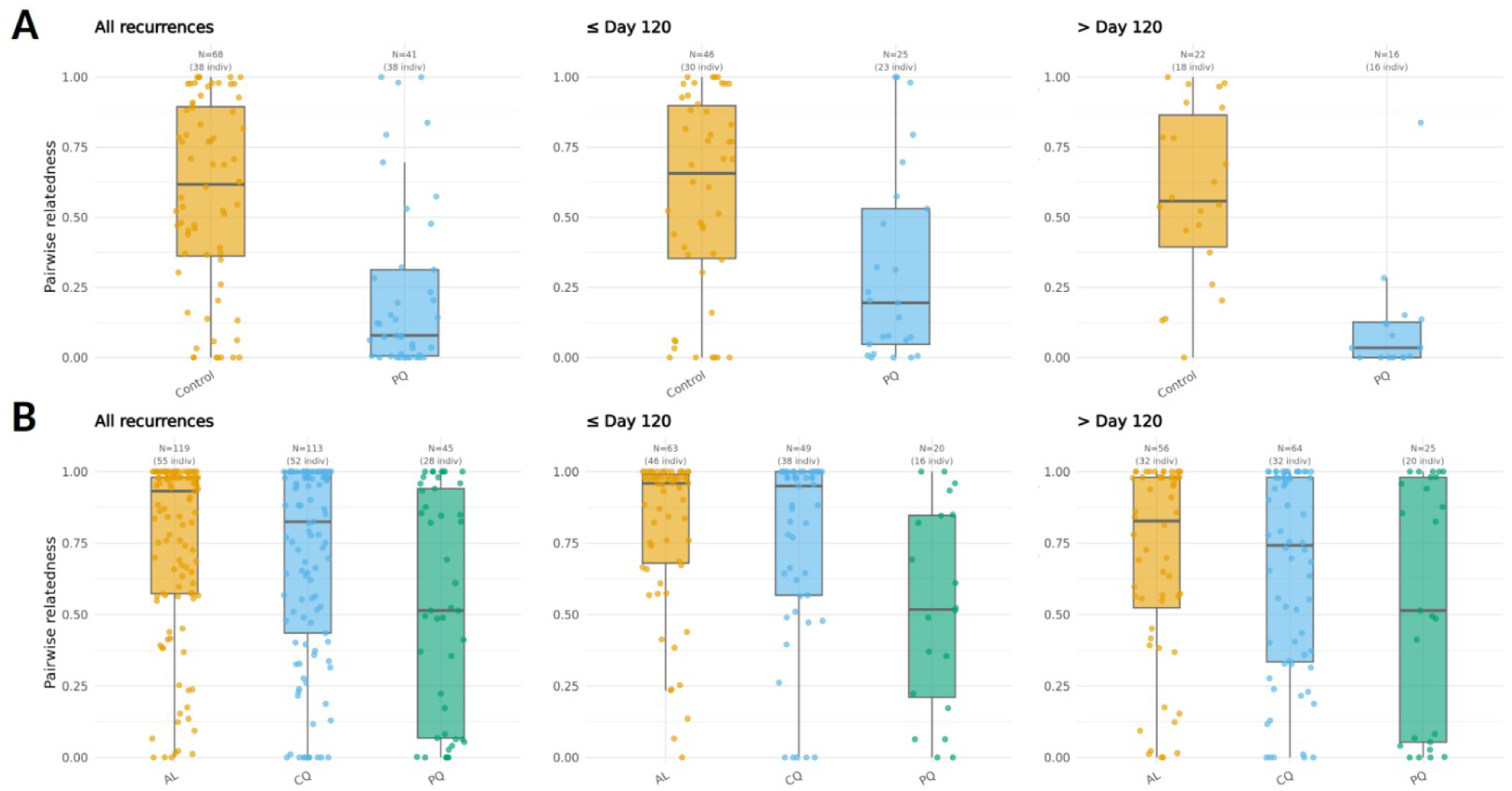
IBD distributions by treatment arm and time to recurrence in IMPROV and OPRA. (A) All IBD distributions between initial infections and all consecutive recurrence pairs grouped by treatment arm in IMPROV (Control Chloroquine [CQ] and Primaquine [PQ]); the same IBD data grouped by recurrences occurring ≤120 days after the initial infection and more than 120 days. (B) All IBD distributions between initial infections and all consecutive recurrence pairs grouped by treatment arm in OPRA (Artemether-Lumefantrine [AL], Chloroquine [CQ], and Primaquine [PQ]); the same IBD data grouped by recurrences occurring ≤120 days after the initial infection and more than 120 days. Each boxplot presents the median and interquartile range (IQR; 25th to 75th percentile), with the Tukey convention whiskers extending to 1.5 x IQR range.

When restricting within-host recurrences to those that occurred prior to day 120, in-line with the typical relapse period of approximately 4 months for Ethiopian *P. vivax* infections, mean IBD in the IMPROV Control arm was estimated to be 58% (N = 46, 30 individuals; IQR 35.3 – 89.8%), compared to 31% (N = 25, 23 individuals; IQR 4.7 – 53.0%) in the PQ arms^19,21^ (**Supplementary Figure S5**). When assessing recurrences beyond 120 days, mean IBD was 58% in the control arm (N = 22, 18 individuals; IQR 39.4 – 86.5%), 28% in the 7-day arm (N = 10; 10 individuals; IQR 8.9 – 51.9%), and 11% (N = 16, 16 individuals; IQR 0.0 – 12.6%) in the 14-day arm. Recurrences before the 120-day threshold, or the theorised relapse window, had a higher mean IBD than those that occurred after day 120 except for the 7-day treatment arm. Estimates of IBD between day 0 and first-recurrence IBD estimates were comparable to the consecutive-pair estimates (Control 59% [N=38]; PQ arms 23% [N=38]).

Within OPRA, mean consecutive-pair IBD was higher in the non-PQ arms, 75% in the AL arm (N = 117; 54 individuals; IQR 57.5 – 98.0%) and 69% in the CQ arm (N = 108; 50 individuals; IQR 40.4 – 98.6%), compared to 55% in the PQ arms (N=43; 27 individuals; IQR 7.5 – 94.9%) (**Supplementary Figure S5**). A Kruskal-Wallis test confirmed a significant difference in relatedness across the three arms overall (H = 9.5, p = 0.0086). A pairwise, Holm-corrected Mann-Whitney U tests showed this was driven by comparisons between the non-PQ arms and the PQ arms (i.e. AL vs. PQ [p = 0.0051]; CQ vs. PQ [p = 0.0452]), while differences between the AL and CQ arms were not significant (p = 0.366). The difference in mean IBD between the PQ and non-PQ arms differed when restricting to monoclonal pairs, with a mean IBD of 85% in the AL arm (N = 69; 39 individuals; IQR 81.3 – 100.0%) and 77% in the CQ arm (N = 60; 36 individuals; IQR 56.7 – 100.0%), compared to 54% in the PQ arms (N = 31; 23 individuals; IQR 6.4 – 97.0%). Day 0 to first-recurrence IBD estimates were comparable to the consecutive-pair estimates (AL = 80% [N = 54]; CQ = 76% [N = 50]; PQ = 50% [N = 28]).

The mean IBD of recurrences occurring before day 120 was 81% in the AL arm (N = 63; 46 individuals; IQR 68.0 – 99.0%) and 75% in the CQ arm (N= 49; 38 individuals; IQR 56.8 – 100.0%), compared to 52% in the PQ arms (N = 20; 16 individuals; IQR 21.0 – 84.7%). For recurrences after day 120, mean IBD was 69% in the AL arm (N = 56; 32 individuals; IQR 52.4 – 98.0%) and 65% in the CQ arm (N = 64; 32 individuals; IQR 33.5 – 98.0%), versus 54% in the PQ arms (N = 25; 20 individuals; IQR 5.4 – 98.0%). There were fewer genotyped individuals in the PQ arms of OPRA who experienced a recurrent episode, with only 16.3% (24/147) experiencing a recurrence, compared to 62.8% (54/86) and 51.6% (49/95) in the AL and CQ arms.

The impact of treatment arm on eMOI per infection episode was assessed to determine if the implementation of radical cure reduced parasite diversity of the hypnozoite reservoir over time but our data did not support this (**Supplementary Figure S6**).

**Supplementary Figure S6:**
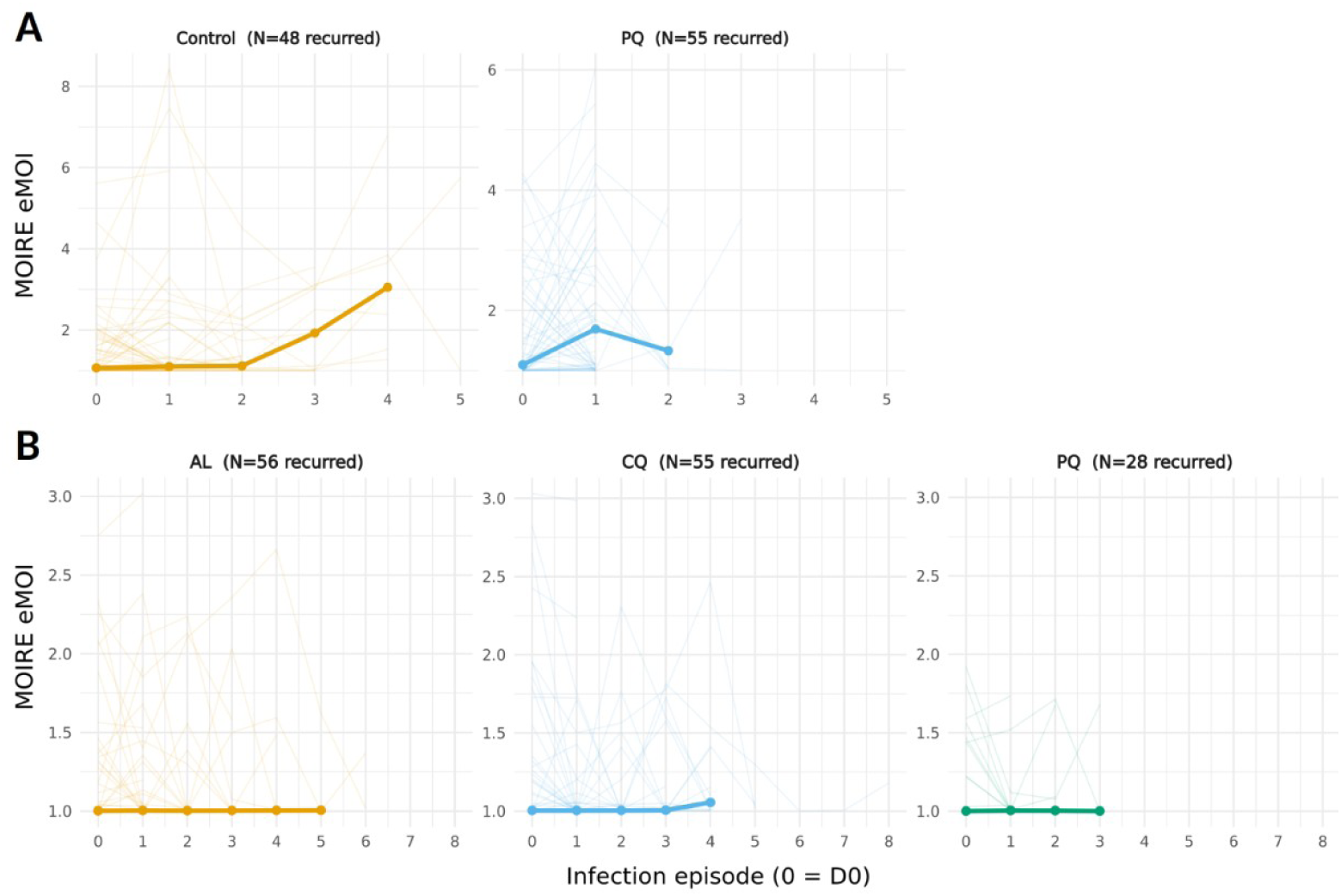
eMOI per recurrence episode by individual; trendline = mean eMOI per episode. (A) IMPROV treatment arms. (B) OPRA trial treatment arms.

### Supplementary Results S3

In IMPROV, monoclonal-only sample sizes were too small in the PQ arms (N = 9 pairs) for a statistically meaningful arm-level comparison. Across all of the IMPROV arms, classification proportions differed significantly between monoclonal-only and polyclonal-involving pairs (χ² = 11.20, df = 2, p = 0.0037), driven substantially by a markedly higher *Pv3Rs* computation failure rate among polyclonal pairs (23.5%; 20/85) than monoclonal pairs (0%; 0/24).

Given the small monoclonal sample size for IMPROV, the description of the monoclonal subset as consistent with the full IMPROV dataset was not supported.

## CODE AVAILABILITY

The *VivaxGEN* ngs-pipeline for retrieving SNP-based variant calls is available on GitHub: github.com/vivaxgen/ngs-pipeline. A GitHub repo for all downstream analyses, *vivaxReC,* is available on GitHub: github.com/aosborne13/vivaxReC/.

## DATA AVAILABILITY

The reads from the genotyped Ethiopian *P. vivax* infections will be made available in the European Nucleotide Archive upon publication.

## AUTHOR CONTRIBUTIONS

A.A.O., R.N.P., and S.A. conceived and designed the study. A. Rai, K.P., and A. Rumaseb contributed to the laboratory coordination and quality assurance. A.A.O, H.T., K.S.H., M.K., and E.S. contributed genotyping data production and informatic support. A.A.O., H.T., and K.S.H. contributed to bioinformatic pipeline development. A.A.O. conducted data analysis. S.R.P., K.T., T.S.D., J.H., A.A., A.W., A.H., A.T., and N.P.D. contributed essential field-based malaria collections and metadata, and guidance on the study design and interpretation.

A.A.O. wrote the first draft of the manuscript, and all authors read and reviewed the manuscript draft. This manuscript has not been accepted or published elsewhere.

## DISCLAIMER

The findings and conclusions in this report are those of the author(s) and do not necessarily represent the official position of the U.S. Centers for Disease Control and Prevention (CDC).

## ACKNOWLEDGEMENTS AND FUNDING

This work was supported, in whole or in part, by the Bill & Melinda Gates Foundation INV-043618 (S.A.). Under the grant conditions of the Foundation, a Creative Commons Attribution 4.0 Generic License has already been assigned to the Author Accepted Manuscript version that might arise from this submission. The work was also supported by the National Health and Medical Research Council of Australia (Investigator Grant 2025377 to S.A.). The OPRA clinical trial was supported by the Bill & Melinda Gates Foundation (OPP1054404 awarded to R.N.P.).

The IMPROV study was funded by the UK Department for International Development, UK Medical Research Council, UK National Institute for Health Research, and the Wellcome Trust through the Joint Global Health Trials Scheme (MR/K007424/1) and the Bill & Melinda Gates Foundation (OPP1054404). We thank the patients who contributed their samples to the study, and the health workers and field teams who assisted with the sample collections. Sequencing was undertaken at the Menzies School of Health Research, Australia, and Australian Genome Research Facility, Australia.

## SUPPLEMENTARY INFORMATION

Supplementary material (Supplementary-Material.xlsx) contains sample metadata, paired sample IDs, trial-specific population-level allele frequencies generated from Day 0 genotyping data, recurrence classification summaries per trial, and individual breakdown of relatedness and recurrence classification probabilities.

